# Establishment & Characterization of a Non-Adherent Insect Cell Line for Cultivated Meat

**DOI:** 10.1101/2024.10.18.618906

**Authors:** Sophia M. Letcher, Olivia P. Calkins, Halla J. Clausi, Aidan McCreary, Barry A. Trimmer, David L. Kaplan

## Abstract

This study presents a blueprint for developing, scaling, and analyzing novel insect cell lines for food. The large-scale production of cultivated meat requires the development and analysis of cell lines that are simple to grow and easy to scale. Insect cells may be a favorable cell source due to their robust growth properties, adaptability to different culture conditions, and resiliency in culture. Cells were isolated from Tobacco hornworm (*Manduca sexta)* embryos and subsequently adapted to single-cell suspension culture in animal-free growth media. Cells were able to reach relatively high cell densities of over 20 million cells per mL in shake flasks. Cell growth data is presented in various culture vessels and spent media analysis was performed to better understand cell metabolic processes. Finally, a preliminary nutritional profile consisting of proximate, amino acid, mineral, and fatty acid analysis is reported.

**Graphical Abstract:** 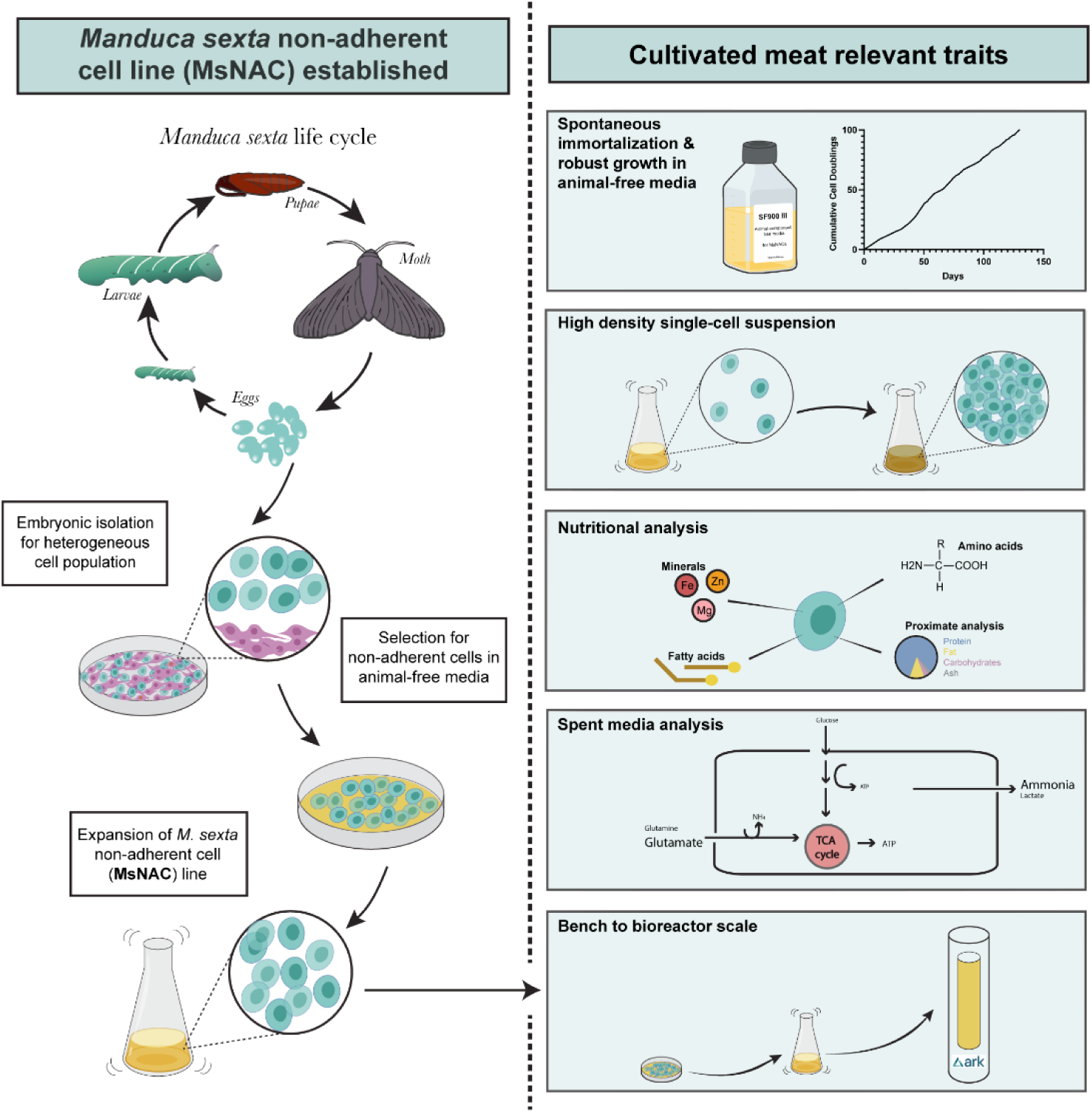

## 1. Introduction

Cultivated meat (CM) is an emerging technology dedicated to producing meat from cell culture instead of whole animals. *Entomoculture*, or culturing insect cells for food, may be a promising alternative to “traditional” livestock cell sources for CM. This is because insect cells are relatively adaptable and resilient in culture, leading to improved scalability at a lower cost. However, studies on insect cell culture bioproduction systems have primarily analyzed cells in the context of generating biologicals (e.g., recombinant proteins, viruses). The present study builds upon previous entomoculture proof-of-concept studies in an effort to demonstrate the feasibility of insect cells as an ingredient in CM products.

### 1.1 Cultivated meat

Growing evidence has shown that traditional animal agriculture is replete with environmental, animal welfare, and human health concerns (Godfray et al., 2018). To mitigate these issues, researchers are investigating how to produce traditionally animal-derived products from cell culture. To date, the focus for the field has been on cells cultured from livestock or seafood species to create cultivated meat (CM) or seafood products. Since the field’s inception approximately 10 years ago, many scientific and regulatory achievements have been accomplished. Notably, in 2023, two companies received permission from the U.S. Food and Drug Administration (FDA) and the U.S. Department of Agriculture (USDA) to commercialize cultivated chicken products made with embryonic chicken fibroblasts, signifying a major milestone for CM production in the United States (Qiu, 2023). In addition to chicken, companies and academic researchers have been investigating bovine, porcine, and piscine cell lines for CM products (Messmer et al., 2023; Pasitka et al., 2023; Saad et al., 2022; Stout et al., 2023a).

While significant progress has been made in developing these cell lines within the constraints of CM (animal-component free cell culture media, easy transition to large-scale culture, favorable nutritional and sensory properties), widescale production remains limited by technical challenges. For example, animal-component free media is available for some cell types but can slow growth relative to fetal bovine serum (FBS)-containing media (Stout et al., 2022), microcarriers are often required for suspension culture (adding complexity and cost) (Bodiou et al., 2020; Hanga et al., 2021), and achieving high cell densities in single-cell suspension is often contingent upon excessive growth media usage (Pasitka et al., 2023).

Insect cells are promising alternatives. Compared to livestock, insects have had incredible evolutionary success, making up the largest and most diverse group of organisms in the animal kingdom (Chapman, 2009). They are capable of surviving in extreme environments and are tolerant of thermal, osmotic and toxic stresses (Law et al., 1992). Many features that make insects resilient and adaptable translate to the cells themselves, which are known for their robust growth, ability to easily scale, and adaptability to different environmental and growth media conditions (Drugmand et al., 2012; Rubio et al., 2019b). The goal of this project was to establish an isolation procedure to develop a new insect cell line that is scalable, easily adaptable to animal-free media and generates sufficient biomass to assess nutritional properties. This approach is intended to provide a framework for future research into other insect species and to further explore the potential of large-scale insect cell culture for consumption. Although the final use-case for entomoculture cell lines compared with traditional biotechnology insect cell culture are different (food ingredients vs. biological production), many of the reasons insect cells are favorable for CM were previously established in adjacent industries.

### 1.2 Insect cells in biotechnology

Historically, motivation for insect cell cultivation has been to produce biologicals, mainly recombinant proteins and biopesticides. As interest grew in using insect cell culture over the past century, focus turned to developing fast-growing cells in large-scale culture. In general, insect cells are favorable for scale-up because they: (1) typically grow well in single-cell suspension and reach higher cell densities than mammalian cells, (2) are easily adaptable to serum- and animal component-free media, thus reducing scale-up costs and batch-to-batch variability of animal components, and (3) grow at near-room temperature and without CO

_s_upplementation, simplifying culture conditions (Drugmand et al., 2012).

The most commonly used insect cell lines in biotechnology are Sf9 cells, isolated from ovarian tissue of the fall armyworm (*Spodoptera frugiperda*) and High-Five cells, isolated from ovarian tissue of the cabbage looper (*Trichoplusia ni*) (Drugmand et al., 2012). Commercial use of insect cells for human therapeutics includes FluBlock (a flu vaccine produced in Sf9 cells) (Cox and Hollister, 2009), Cervarix (a cervical cancer vaccine produced in High-Five cells) (Schiller et al., 2008), and NovaVax (a COVID-19 vaccine that uses Sf9 cells to produce a recombinant SARS-CoV-2 spike protein) (Pijlman et al., 2020; World Health Organization, 2021). These industries provide a precedent for the scalability of insect cell culture and a solid foundation for further research into using insect cells for food.

### 1.3 Entomoculture

In 2007, researchers at Wageningen University proposed using insect cells cultivated in bioreactors to generate edible protein. The rationale behind this idea was that whole insects are nutritionally favorable with respect to their mineral and digestible protein content, and *in vitro* growth offers a high level of control and customization (Verkerk et al., 2007). As CM gained traction in the past decade, the idea of growing insect cells for consumption (*entomoculture*) resurfaced, again motivated by the fact that infrastructure already exists for large-scale insect cell culture, and compared with “traditional” livestock species, insect cells are extremely adaptable and resilient in culture (Rubio et al., 2019b).

Entomoculture does not necessarily aim to recreate edible insect food products *in vitro –* rather, it aims to use insect cells as the animal cell ingredient in a variety of cultivated meat products, providing the sensory and nutritional advantages of animal meat with a potentially easier and more cost-effective path to scale-up than conventional livestock species. The advantageous qualities of insect cells that render them suitable for production of biologicals also position them as an efficient system for generating biomass directly from the cells themselves. Incubation systems can be simplified and facility costs lowered by room-temperature growth without CO_2,_ animal components can be removed from biomass production with easy adaptation to animal-free media formulations, and the lack of contact inhibition can help to reach higher cell densities (Letcher et al., 2022; Rubio et al., 2019a). Insect cells also grow well in the absence of growth factors such as insulin, transferrin, fibroblast growth factor, and transforming growth factor beta, which are typically used for mammalian cell culture and contribute to the high cost for large-scale CM (Humbird, 2021; Rubio et al., 2019b). Despite the potential for insect cells for CM, previous research into developing and studying insect cells in a food context has been extremely limited.

### 1.4 Previous work

Research in our laboratory has shown proof-of-concept data supporting the use of insect cells for CM, however cells have not been analyzed beyond small bench-scale. For example, a *Drosophila melanogaster* (fruit fly) muscle cell line was adapted to serum-free media and formed differentiated muscle fibers in chitosan films and sponges (Rubio et al., 2019a). In this study, cells were adapted to single-cell suspension using dextran sulfate to reduce cell aggregation in bench-scale shake flasks at low cell densities (Rubio et al., 2019a). Embryonic cells from *Manduca sexta* (tobacco hornworm caterpillar) have also been explored for cultivated fat, accumulating lipids upon treatment with a soybean oil emulsion, with a fatty acid breakdown similar to whole caterpillars (Letcher et al., 2022). A preliminary technoeconomic assessment has also been performed, which projects that insect cell lines cultured in the context of CM have significantly lower media, oxygen consumption, and utility costs compared with mammalian cells, resulting in lower production costs per kilogram (Ashizawa et al., 2022).

Although insect cells have been used in biotechnology for biological production for decades, there are gaps in strategies to generate new cell lines optimized for food-related goals. Here, we demonstrate the generation of an insect cell line from *M. sexta* embryos, with cell line development and analysis specifically tailored toward the goals of CM. *M. sexta* was selected as it is a relatively well-characterized lepidopteran model organism. *M. sexta*’s developmental cycle includes an embryonic, larval (caterpillar), pupal, and adult (moth) stage (Dorn et al., 1987; Gershman et al., 2021). Cells were isolated from embryos because previous research has shown robust embryonic cell growth, presenting an opportunity to select for desirable traits (e.g., growth in suspension and animal-free culture media) (Baryshyan et al., 2012; Letcher et al., 2022). Once a cell line was established that showed robust growth in animal-free culture media, traits relevant for CM production and consumption were explored, including high-density growth in single-cell suspension, spent media analysis, and nutritional profiling.

## 2. Materials & Methods

### Cell Isolation

*M. sexta* non-adherent cells (MsNACs) were isolated as described previously (Baryshyan et al., 2012) with some alterations. Briefly, *M. sexta* eggs were staged at 19-22 hours (stage three of development (Dorn et al., 1987)), sterilized with 50% sodium hypochlorite solution for 5 minutes, rinsed with growth media and lysed using a Dounce homogenizer. Homogenate was filtered through a 70-micron filter, centrifuged at 40 *xg* for 10 minutes to remove debris, and the supernatant was transferred to a fresh tube and centrifuged for 10 minutes at 380 *xg*. The top layer of the supernatant (1 mL) was transferred to a 12.5 cm^2^ T-flask that was filled completely with growth media and incubated at 27°C for 12 days. After 12 days, media was refreshed, and a 50% media exchange was performed twice over two weeks. Non-adherent cells were routinely subcultured at 90% confluency by lightly pipetting media at the bottom of the flask to detach cells and seeded at 100,000 cells/cm^2^. Initial growth media was Shields and Sang M3 media (S8398, Sigma-Aldrich, St. Louis MO) supplemented with 0.5 g/L potassium bicarbonate, 2.5 g/L bactopeptone, 1 g/L yeast extract and 10 (vol./vol.)% heat inactivated fetal bovine serum. Cells were transitioned to SF900 III (12658019, ThermoFisher, Waltham, MA) after passage 4 (see Serum-Free Media Adaptation, section 2.2). 1X Antibiotic-Antimycotic (ThermoFisher 15240096) was used during preliminary culture but was removed after the third passage.

### Serum-Free Media Adaptation/Nonadherent Cell Selection

After 4 passages in growth media containing 10% fetal bovine serum, cells were gradually adapted to SF900 III animal component-free media (ThermoFisher 12658019). Cells were passaged into 75% serum-containing and 25% serum-free media at 100,000 cells/cm^2^. Every 5 days, non-adherent cells were collected and seeded in a new flask with 25% reduced serum until cells were cultured in 100% SF900 III animal component-free media. For 6 months, cells were passaged each week by collecting nonadherent/lightly adherent cells by gently pipetting media over the bottom of the culture flask, removing the media containing floating cells, and diluting the media and cell suspension 1:3 into a fresh T-flask. After 6 months, cells were transferred to 125 mL shake flasks and maintained in suspension for the duration of the study unless otherwise noted.

### Cryopreservation

MsNACs were expanded in cell culture flasks and harvested after reaching 90% confluency. Cells were centrifuged at 380 *xg* for 5 minutes and resuspended at 10E6 cells/mL in freezing media (45% conditioned media, 45% fresh media and 10% dimethyl sulfoxide). Cells were transferred to cryovials and frozen in a cryobowl at 80°C overnight before relocation to long-term liquid nitrogen storage. Cell viability and doubling time was determined from four different passages with an automated cell counter (NC-200™, Chemometec) after thawing cryopreserved cells.

Each vial of cryopreserved cells was allowed to recover for 4-5 days after thawing in a T25 flask. After one week, 100,000 viable cells/cm^2^ were passaged into a fresh flask and cell viability and doubling time was determined with an automated cell counter (NC-200™, Chemometec) after 5 days of growth. This process was repeated for four separate cryopreserved vials (P34, P36, P41, P42) and measurements were compared to doubling time/viability recorded prior to cryopreservation.

### Growth Curves

MsNACs were seeded at 500,000 cells/mL in 30 mL working volume in 125-mL shake flasks (ThermoFisher 4113-0125) in triplicate. For highest achievable cell density studies, 120 µL cell suspension was removed at 24-hour intervals and cell count and viability were determined with an automated cell counter (NC-200™, Chemometec). Once cells reached over 3E6 cells/mL, a 50% media exchange was performed every other day by removing half of the media and cell suspension, centrifuging at 300 *xg* for 5 minutes, aspirating spent media, and resuspending cell pellet with fresh media before adding it back to the flask. For batch growth curves, cells were counted each day without media exchange until growth rates plateaued.

### Bioreactor Runs

Bioreactor experiments were carried out in a 2.4-liter airlift bioreactor. All glass bioreactor components were pre-coated with Sigmacote, (SL2, Sigma-Aldrich) dried overnight, and thoroughly rinsed before use. Passage 30-40 MsNACs were expanded in shake flasks before inoculation into the 2.4-L bioreactor at a viable cell density of 500,000 cells/mL. Culture media was supplemented with 0.5% Antibiotic-Antimycotic (15240096, ThermoFisher Scientific). The bioreactor was operated in batch mode, with media only added as needed to counteract volume lost through evaporation and sampling. The bioreactor was operated at 27°C with an air sparge rate of 0.5 LPM. 1-2 mL of a 1:100 mixture of an antifoam agent and Reverse Osmosis Deionized water was added as needed to minimize foaming levels.

For harvesting cells used in section 2.8, the cell culture was removed in aliquots of 500 mL and spun down at 325xg for 13 minutes. The spent media was aspirated, and the resulting cell pellets were resuspended in 5mL of Phosphate-Buffered Saline to wash, combined into one 50 mL tube, and re-spun down at the same conditions. The Phosphate-Buffered Saline was aspirated, and the resulting pellet was weighed before being stored at -20°C until further analysis. Figure 2 represents cell counts taken with an automated cell counter (NC-200™, Chemometec) at matched timepoints to a parallel 500 mL shake flasks over three passages. Additional measurements (beyond the matched timepoints) were recorded for the 2.4L bioreactor over the three passages and are shown in SI Figure 2.

**Figure 1.**
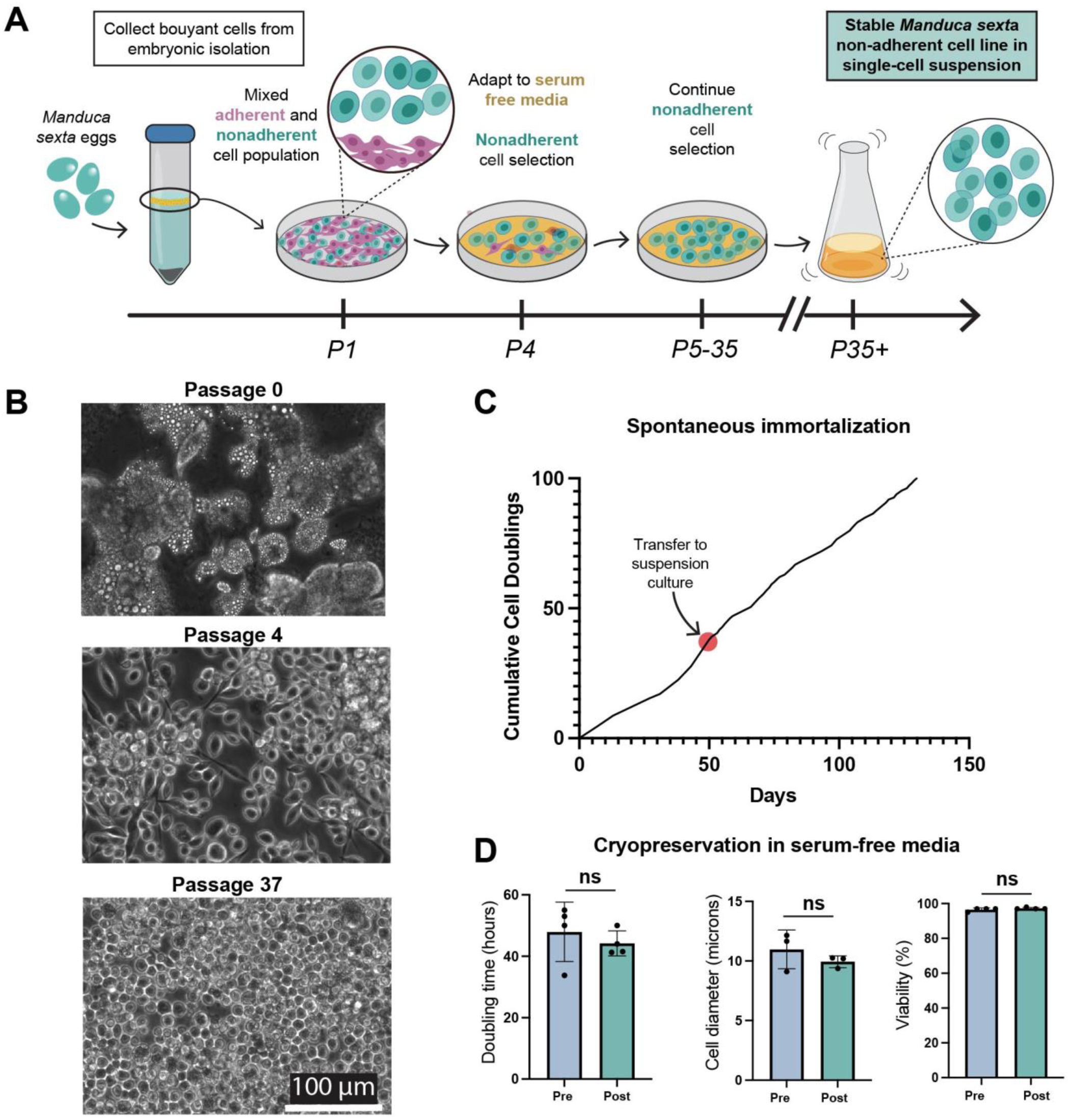
Establishment of spontaneously immortalized embryonic Manduca sexta cell population that proliferates in single-cell suspension in animal-component free media. (A) Schematic of cell line development process. (B) Phase contrast images of cells isolated from buoyant layer at passage 0 (top), passage 4 (middle), and passage 37 (bottom). Scale bar = 100 microns. (C) Tracking of cell doublings up to 100 cumulative cell doublings, representing spontaneous cell immortalization. (D) Analysis of cells before and after cryopreservation; doubling time (left), cell diameter in microns (middle), and viability (right) n=3-4, t-test, ns=not significant.

**Figure 2:**
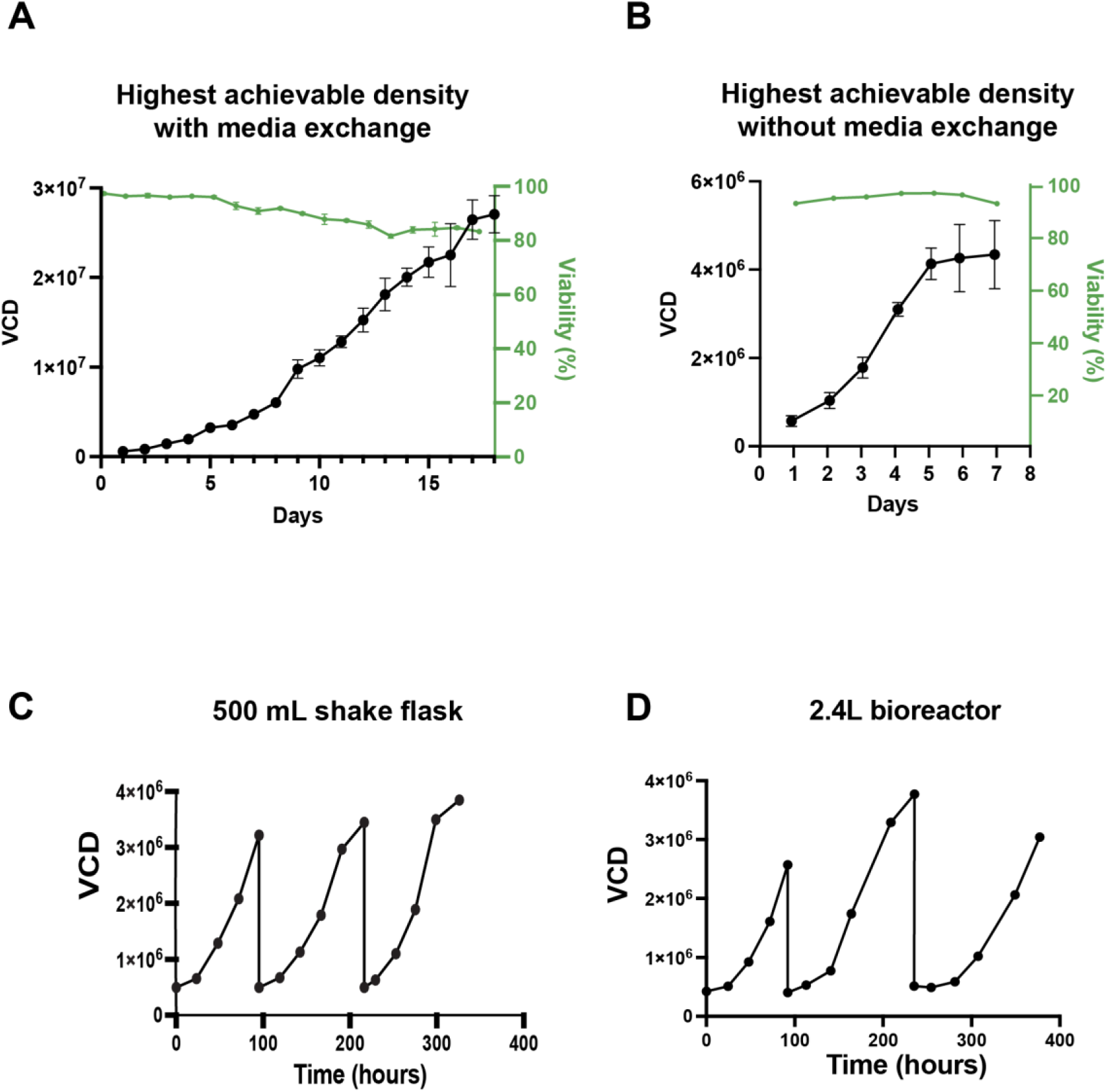
Suspension growth of MsNACs. (A) Highest achievable density of MsNACs with media exchange on left axis, with viability shown on the right axis (n=3). (B) Highest achievable cell density of MsNACs without media exchange on left axis, with viability shown on the right axis (n=3). (C) Growth over three passages in 500 mL shake flasks. (D) Growth over three passages in a 2.4-liter bioreactor.

### Spent Media Analysis

During batch shake flask culture, each day 500 µL was removed from shake flasks. Cell suspension was spun down at 300*xg* and 450 µL cleared supernatant was preserved at -80°C for further analysis using either ViCELL Metaflex bioanalyzer (Beckman Coulter) or Cedex Bio Analyzer (Roche). Throughout the bioreactor runs, samples were taken 2x daily. Glucose and lactate were analyzed using the Vi-CELL Metaflex bioanalyzer (Beckman Coulter) and all other metabolites were measured on a Cedex Bio Analyzer (Roche). All bioreactor metabolite measurements were performed at the time of sampling.

### Ammonia and Lactate Supplementation

Passage 50 MsNACs were expanded in shake flasks before being seeded into 48-well plates to assess growth and viability. Serial dilutions of either ammonium chloride (ThermoFisher, A15000.0I) or sodium lactate (ThermoFisher, L14500.06) were prepared from 100 to 3.13 mM in Sf900 III medium. Cells were seeded at 50,000 cells/cm^2^ in 48-well plates with 5 replicates of each condition. Over 5 days, confluency was monitored with a Celigo Image Cytometer (Nexalom Bioscience). After 7 days, cells were imaged using a phase contrast microscope (BZX-810, Keyence) and viability was analyzed using a LIVE/DEAD stain (ThermoFisher, R37601), and again imaged using a fluorescence microscope (BZX-810, Keyence). Figures 5C and E represent proportions of total cells analyzed that were live and dead. Significance represents an ANOVA performed on the proportion of live cells in each condition.

### Proximate Nutritional Analysis

Three separate 10-15 gram (wet weight) MsNAC samples were collected as described in section 2.5 (“Bioreactor Runs”) above and stored at - 20°C until further analysis. Proximate analysis of MsNAC cell samples was conducted at Eurofins Scientific, Inc. (Eurofins Nutrition Analysis Center, Des Moines, IA). An abbreviated Nutrition Labeling and Education Act (NLEA) analysis was performed with methods established by the Association of Official Analytical Chemists (AOAC). Moisture content was analyzed by forced draft oven (AOAC 925.09), protein by combustion in a protein analyzer (AOAC 990.03, AOAC 992.15), ash by heating at 600°C (AOAC 942.05), and crude fat by acid hydrolysis (AOAC 954.02). Carbohydrate content was calculated by subtracting the sum of ash content, crude fat content, and protein content from 100%. Data is represented as dry weight calculated by proportions of each component as a percent of all measurements except moisture.

### Amino Acid, Fatty Acid, Mineral Analysis

Samples were collected from either MsNACs or DF-1 cells grown in single-cell suspension. MsNACs were seeded at 500,000 cells/mL in 30 mL working volume in 125-mL shake flasks (ThermoFisher 4113-0125) in a shaking incubator set at 27°C with 110 RPM agitation. DF-1 cells were cultured in Dulbecco’s Modified Eagle Medium + 10% Fetal Bovine Serum + 1% antibiotic/antimycotic. Cells were inoculated at 50,000 cells/cm^2^ in a 39°C shaking incubator with 5% CO_2 s_et at 80 RPM. For both cell types, 50 million cells were removed from separate flasks, centrifuged at 300 *xg* for 5 minutes, rinsed with sterile PBS, and cell pellets stored at -20°C until further analysis. Separate samples (*n*=4 for MsNACs, *n*=3 for DF-1) were collected for amino/fatty acid and for metal analysis. Food composition assays were performed at The Metabolomics Innovation Centre (TMIC) at the University of Alberta. Briefly, cells were lysed with 50% methanol and 50% water and four rounds of freeze-thaw in liquid nitrogen. Each sample was centrifuged at 16000 *xg* for 10 minutes, and the supernatant used for Liquid Chromatography with Tandem Mass Spectrometry (LC-MS/MS) analysis. A targeted quantitative metabolomics approach was used as previously described (Zheng et al., 2021). A direct injection mass spectrometry with a reverse-phase LC-MS/MS custom assay was combined with an ABSciex 4000 QTrap (Applied Biosystems/MDS Sciex) mass spectrometer. Data was analyzed using Analyst 1.6.2. Metabolites were quantified using isotope-labeled internal standards. For metal analysis, Inductively Coupled Plasma Mass Spectrometry (ICP-MS) was performed on a NexION 350x ICP–MS (Perkin-Elmer, Woodbridge, ON, Canada) using previously described methods (Foroutan et al., 2020, 2019). One 20-gram sample was sent to Eurofins for Amino Acid analysis via acid hydrolysis (Eurofins code QQ176). Results from Eurofins were reported as percentages, and SI Figure 4C represents percentage of each amino acid of total analyzed amino acids.

### DF-1 Comparisons

Protein concentrations of MsNAC and DF-1 cells were compared via a Bicinchoninic Acid Kit (BCA) (A55864, Thermo Scientific). MsNAC and DF-1 cell samples were centrifuged at 600*xg* for 5 minutes to isolate the cell pellet, rinsed with Phosphate Buffered Saline, and resuspended and lysed in Radioimmunoprecipitation Assay Buffer (RIPA buffer). Samples were then centrifuged at 14000*xg* for 10 min and supernatant was transferred into new tubes for each condition. BCA standards, using RIPA as diluent, and BCA working reagent were prepared as outlined in the Thermo Scientific Pierce™ BCA Protein Assay Kit protocol. Standards, samples, and working reagents were transferred into a 96-well plate at a sample to working reagent ratio of 1:8. The well plate was incubated at 37°C for 30 minutes and absorbance was measured at 562 nm on a Synergy H1 microplate reader (BioTek Instruments, Winooski, VT, USA). DF-1 and MsNACs were imaged via light microscope at 40x and subsequently analyzed for area and diameter using ImageJ.

### Statistical Analysis

Statistical analysis was performed with GraphPad Prism Version 9 software (San Diego, CA, USA). All experiments were performed in triplicate unless otherwise noted in figure legends. Unpaired t-tests were performed for comparing means between two samples (i.e., the mean doubling time, cell diameter, and viability collected from 3-4 replicates of cells from different passages before vs. after cryopreservation in Figure 1, and the mean concentrations of metals (in micromolar) from 3-4 separate flasks of MsNACs compared with DF-1 cells in Figure 4). For comparisons of more than two samples, a one-way Analysis of Variance (ANOVA) was performed to compare multiple groups with a single factor (e.g., cell viability with various ammonia or lactate concentrations in Figure 3). A two-way ANOVA was performed for experiments involving two independent variables (e.g., proportions of essential amino acids compared between MsNACs and DF-1 cells in Figure 4). A mixed effects ANOVA was performed for experiments with both independent variables and repeated measures (e.g., percent confluency over multiple days with various lactate or ammonia concentrations). Multiple comparisons were analyzed via Dunnet’s post-hoc test unless otherwise noted in figure legend. P-values < 0.05 were considered significant. Errors represent mean ± standard deviation.

**Figure 3.**
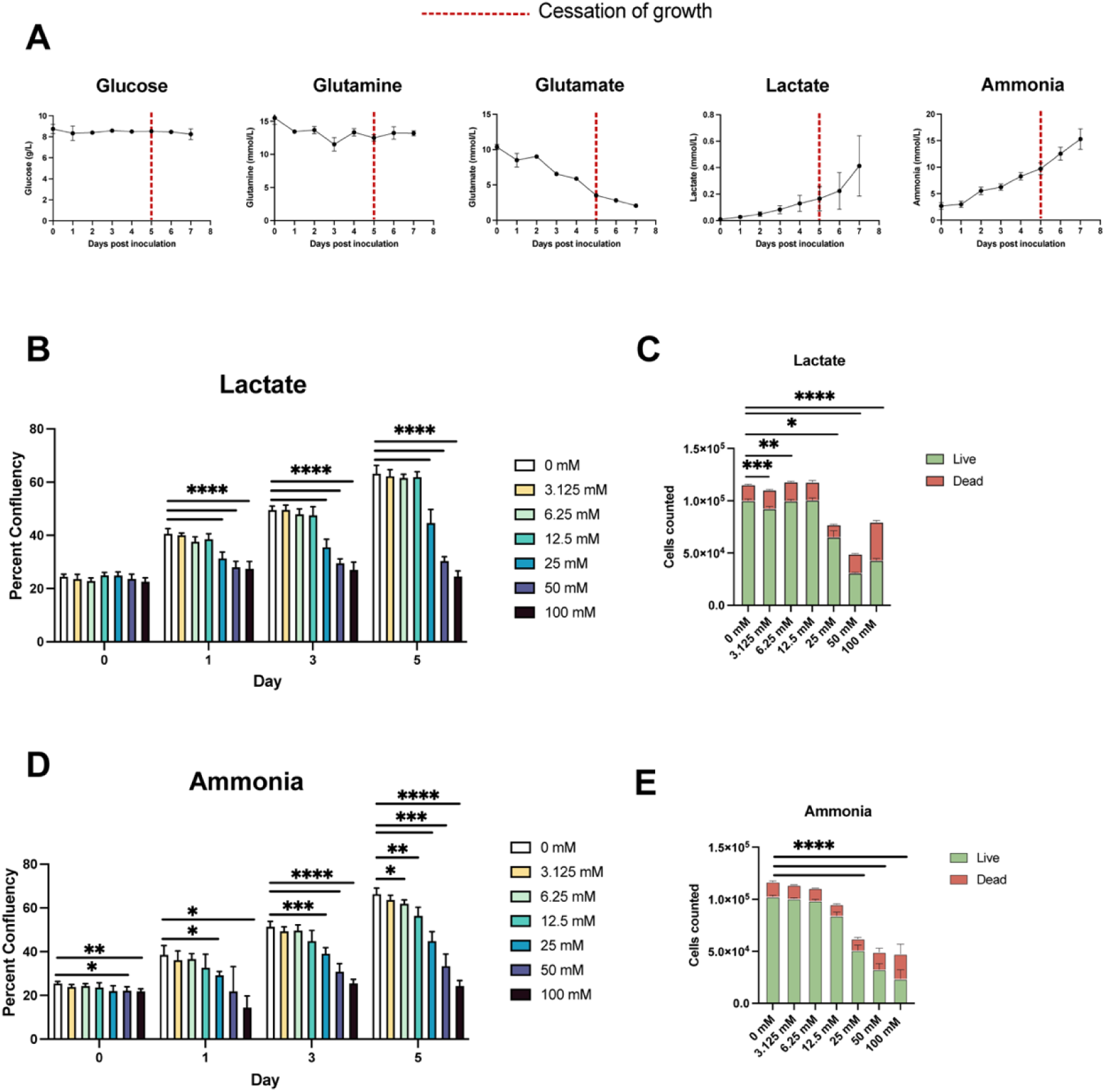
Spent media analysis and inhibitory concentrations of lactate and ammonia. (A) Spent media analysis for ammonia, glutamate, l-glutamine, glucose and lactate for cells grown in Figure 2B, with red line indicating timepoint of growth cessation (n=3 flasks) (B) Growth over 5 days as measured by confluency in 48-well plates for MsNACs growth with increasing concentrations of sodium lactate. Mixed effects ANOVA with Dunnett’s multiple comparisons test (n =5). (C) Proportion of live and dead cells as measured by calcien AM (live) and propidium iodide (dead) after 7 days of growth with increasing concentrations of sodium lactate. 1-way ANOVA with Dunnett’s multiple comparisons test (n=3) (D) Growth over 5 days as measured by confluency in 48-well plates for MsNACs growth with increasing concentrations of ammonium chloride. Mixed effects ANOVA with Dunnett’s multiple comparisons test (n =5). (E) Proportion of live and dead cells as measured by calcien AM (live) and propidium iodide after 7 days of growth with increasing concentrations of ammonium chloride. 1-way ANOVA with Dunnett’s multiple comparisons test (n=3). *<0.05, **<0.01, ***<0.001, ****<0.0001.

**Figure 4:**
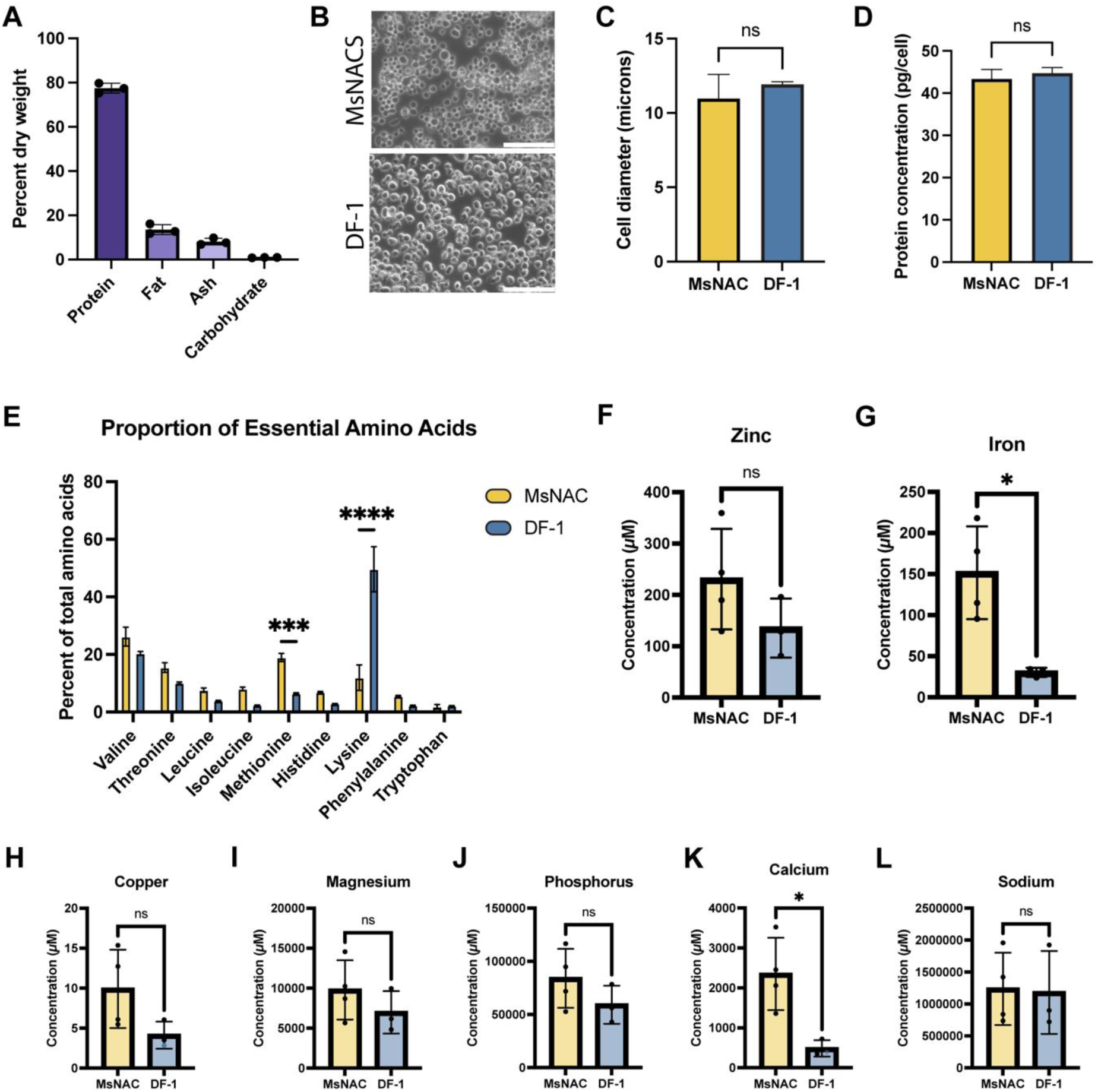
Nutritional analysis of MsNACs. (A) Dry weight breakdown as analyzed via combustion of 3 separate 15-20 gram samples of MsNACs. (B) phase contrast images of MsNACs (top) and DF-1 cells (bottom) SB = 100 microns. (C) Cell size comparison between MsNACs and DF-1 cells as analyzed by ImageJ (n=3 fields of view, 30-80 cells per field of view). (D) Protein concentration as measured in picograms per cell via BCA assay (n=3). (E) Proportion of each essential amino acid as a percent of all amino acids analyzed via UPLC-MRM/MR of MsNAC and DF-1 cells. n=3-4, 2-way ANOVA with Sidak’s multiple comparisons test ***<0.001, *****<0.0001. (F-L) comparison of MsNAC (yellow) and DF-1 (blue) metals via ICP-MS reported in micromolar. t-test, *<0.05 ns = not significant.

## 3. Results

### 3.1 Primary Culture: MsNAC culture yielded cells that proliferate in animal-origin free media

Cells were isolated from *M. sexta* embryos as shown in Figure 1. Although a range of cell morphologies were observed from the cell pellet after centrifugation (SI Figure 1A), the initial population from the floating layer appeared to primarily consist of lipid-filled cells (Figure 1B). After media refreshment, a relatively homogenous and rapidly proliferating cell population emerged and was passaged (Figure 1B). Cells could also be expanded from the pellet, but cells from the floating layer proliferated faster and a larger proportion were nonadherent, and thus were selected for further expansion. Because the goal of this study was to generate cells that could proliferate in single-cell suspension, only non-adherent or lightly adherent cells were expanded. Fetal bovine serum and antimicrobials were removed after four passages. Interestingly, once serum was removed cells adhered more tightly to the tissue culture plastic and an elongated morphology was observed (SI Figure 1B). However, further selection for non-adherent cells over multiple passages in Sf900 III media resulted once again in a more rounded phenotype (Figure 1B).

Selection for non-adherent cells in static culture in Sf900 III media was continued for approximately 6 months. At passage 25, cell morphology, viability, and growth rate appeared consistent, and cells were named “*Manduca sexta* Non-Adherent Cells,” or “MsNACs.” Cell expansion continued, and consistent growth rates were observed as cells reached over 100 doublings, indicating that MsNACs had spontaneously immortalized (Figure 1C). After approximately 40 doublings, cells were transferred to suspension culture. Cells continued to grow at consistent rates past 100 doublings and have been maintained as a continuous culture in suspension for over 1.5 years with an average doubling time of 33.5 +/- 6.3 hours between passage 80-95 (SI Table 1).

Growth rate and viability before and after cryopreservation was analyzed, as cell banking is essential for large-scale cell expansion. As the high protein content in fetal bovine serum aids in protecting cells during cryopreservation, serum-free cryopreservation can be challenging. Although cells exhibited poor (<50%) viability upon initial thawing (data not shown), after one passage and approximately one week of recovery the doubling times and viability did not significantly differ from pre-cryopreservation numbers (Figure 1D). Of note, doubling times represent total cell growth over five days in static culture; log phase doubling time decreased to ∼30 hours in both conditions. Morphology also appears unchanged after thawing (SI Figure 1D), and cell size did not change (Figure 1). In addition, cells were confirmed to be mycoplasma free across multiple passages (SI Figure 1E). Finally, species validation was performed to confirm the expanded cells were indeed derived from the original isolation. Cells from three separate passages were confirmed to be 99.25 +/- 0.29% similar to published *M. sexta* COI sequences, with less similarity to the related species *Manduca quinquemaculata* (the tomato hornworm), and significantly different from other commonly studied lepidoptera species, *Bombyx mori* and *Spodoptera frugiperda* (97.39 +/- 0.29%, 89 +/- 0.28%, and 89 +/- 1.85%, respectively, SI Figure 1C).

### 3.2 MsNACs proliferate in shake flask culture and reached densities over 20E6 cells/mL

MsNACs reached over 25E6 cells/mL in shake flasks with media exchange before growth rates slowed dramatically (Figure 2A). Although cell viability dropped below 90% at 8-10E6 cells/mL, it never decreased below 80% for the duration of the study (Figure 3A). Similar trends were observed in cells at higher passages (P50) after cryopreservation, although cell viability at densities over approximately 10E6 cells/mL were decreased (SI Figure 2A). In batch culture without media exchange, cells reached an average of 4.34 +/- 0.77E6 cells/mL over 7 days, with viability remaining above 92% (Figure 2B). Optimization of biomass production was also explored by comparing media exchange (removal and replacement of 50% of the media every other day) with fed-batch (adding 5 mL of fresh media every other day to a 125-mL shake flask culture without removing spent media). Although viable cell densities remained similar, after 8 days the fed-batch had significantly more total cells as there was a greater total volume with the same cells/mL (SI Fig 2D). This is a favorable approach, as fed-batch offers a simpler feeding paradigm and uses less media overall. Two other commercial insect animal-origin free media were explored: ExpiSF™ CDM (Thermofisher A3767801) and ESF AF™ (VWR 101419-848). Cells were gradually adapted to the new media formulations over 4 passages with 25% increases in the new media each passage. At 75% new medium and 25% Sf900 III media, cells were observed to have unhealthy morphology in ESF AF™ media and ceased growth (data not shown). In ExpiSF™ CDM, cells had slightly longer lag phases compared with Sf900 III and reached similar cell densities during starvation, thus Sf900 III media was selected for scale-up and spent media studies (Supplemental Figure 2E).

### 3.3 MsNACs were scaled to a 2.4-liter bioreactor

Once consistent cell growth was observed in small-scale shake flasks, cells were grown in benchtop pneumatic bioreactors to determine feasibility of a MsNAC at-scale production process. Over the three passages, cell viability in the 2.4L bioreactor was generally lower compared to 500-mL shake flasks but did not drop below 80% (results not shown), while cell densities were similar between the two vessels over three passages (Figure 2C,D). Doubling times in the 2.4L were 44.32 +/- 8.77 hours from inoculation to passage and dropped to 23.6 +/- 7.9 during the fastest period of growth (on either day 2 or 3 of growth, SI Table 1). MsNACs were also successfully grown in a 10-L bioreactor, reaching a peak density of approximately 2E6 cells/mL with slower than expected growth (SI Fig 2B). With further optimization of culture conditions at this larger scale, more efficient cell growth would be anticipated.

### 3.4 Spent media analysis offers insight into MsNAC metabolism

Although cells survive and grow at densities above 20E6 cells/mL, without media exchange cell density typically cannot reach above 4E6 cells/mL in SF900 III media (Figure 2). Spent media from cells grown in Figure 2B was analyzed to understand why growth ceased. Throughout the course of 7 days of batch culture (with growth slowing around day 5), glutamate was largely exhausted, but interestingly glucose was not consumed, and glutamine fluctuated but remained above 10 mmol at day 7 (Figure 3A). Metabolic byproducts were also analyzed: lactate was produced at extremely low rates (less than 1 mmol), and ammonia was consistently produced and reached over 15 mmol over 7 days (Figure 3A). Similar trends were observed in the 2.4L bioreactor, indicating that cell metabolism was consistent regardless of culture vessel (SI Figure 3B). Dissolved oxygen remained relatively stable, but carbon dioxide and the pH of the spent media increased over the 7-day period (Supplemental figure 3A).

Because metabolite buildup likely inhibits cell growth, the effects of lactate and ammonia were tested by adding increasing concentrations of either sodium lactate or ammonium chloride to cells in fresh media and observing cell growth. MsNACs appear to be unaffected by sodium lactate up to 12.5 mM (although the proportion of live cells decreased slightly), but growth was drastically decreased above 25 mM (Figure 3B,C, SI Figure 3E). Since lactate production is normally below 1 mM, it is unlikely that it contributes to the slowed growth. In contrast, ammonium chloride added to cells had a detrimental effect within the range of ammonia produced by cells in Figure 3A. After five days exposure to concentrations above 6.25 mM growth was slowed, although the most dramatic effects were seen above 25 mM (Figure 3D,E). Notably these effects were not due to changes in pH, which remained between 6.09-6.22 in all conditions (SI Figure 3C).

### 3.5 Analysis of MsNAC Nutritional Profile

#### 3.5.1 Proximate analysis, amino acid, fatty acid, minerals

Preliminary nutritional analyses were performed on MsNACs grown in single-cell suspension in animal-component free medium in 2.4 and 10-liter bioreactors. Dry weight calculations showed MsNACs consisted of 77.47 +/-2.2% protein, 13.53 +/- 2.27% fat, 8.1 +/- 1.51% ash, and 0.92 +/- 0.1% carbohydrate (Figure 4A). MsNACs had all nine essential amino acids based on UPLC-MRM/MR analysis, with individual breakdowns (µM) shown in SI Table 2. Metal analysis was also performed via ICP-MS, and results are reported (µM) in SI Table 2. Heavy metals (lead, arsenic, and cadmium) were not detected.

#### 3.5.2 Comparisons to embryonic chicken fibroblasts

To place these nutritional data in context, MsNACs were compared to embryonic chicken fibroblasts (DF-1 cells). Spontaneously immortalized embryonic chicken fibroblasts are currently the only CM generated cells approved for consumption through the Food and Drug Administration (FDA) and the United States Department of Agriculture (USDA), who jointly regulate CM in the United States (Failla et al., 2023). Data for the dry weight composition and amino acid proportions were obtained from safety documentation published by the California-based cultivated meat companies Good Meat and UPSIDE, who both use embryonic chicken fibroblasts for their cultivated chicken products. Data was also obtained from DF-1 cells cultured in single-cell suspension in-house for the metal analysis and the amino acid analysis (Figure 4E-L). Breakdown of dry weight protein, fat, ash, and carbohydrates were similar between MsNAC and the embryonic chicken fibroblasts from GOOD meat and UPSIDE; MsNACs had significantly higher carbohydrates compared with GOOD meat, but significant differences were also seen between UPSIDE and GOOD protein and ash composition (SI Fig 4A). MsNACs had significantly decreased proportions of lysine, threonine, and valine compared with DF-1 cells (SI Fig 4B). When comparing normalized proportions of essential amino acids, MsNACs had significantly more methionine and significantly less lysine compared with DF-1 cells (Figure 4E). All other amino acids had no significant differences (Figure 4E). One 20-gram wet cell weight sample was also analyzed by acid hydrolysis, with percentages of amino acids shown in SI Fig 4C. The relative amount of each amino acid was similar to embryonic chicken fibroblasts and conventional chicken as reported in safety dossiers from two separate CM companies (SI Figure 4C). There were no significant differences in zinc, magnesium, copper, sodium, and phosphorous concentrations between cell types, however MsNACs had significantly more iron and calcium compared with DF-1s (SI Table 4, 4F-L). Finally, fatty acids were analyzed via LC-MS/MS and compared between MsNACs and DF-1 cells (SI Fig 4D, SI Tables 5-8). Results are broken down into free fatty acids, lysophosphatidylcholines (LysoPC a), and phosphatidylcholine diacyl (PC aa). There are minimal differences between DF-1 and MsNACs on the fatty acid level, although MsNACs have significantly higher C2 (Acetic Acid), PC aa C32:2 and PC aa 34:2. Besides PC aa 32:2 and PC aa 34:2, DF-1s have significantly more of many PC aas.

## 4. Discussion

Widespread adoption of CM is contingent upon the generation of scalable, cost-effective, meat-relevant cell lines. Here, we present a relatively simple protocol to develop lepidopteran cell lines from embryos that can be grown in animal-free culture media and easily adapted to high-density single-cell suspension culture. Although insect cells have been grown for decades to produce biologicals, cell line development and analysis has previously been constricted to basic growth kinetics and recombinant protein/viral output (Beas-Catena et al., 2011; Hashimoto et al., 2010; Stavroulakis et al., 1991). In contrast, the *Manduca sexta* non-adherent cells (MsNACs) generated here were analyzed with the goal of biomass production for consumption. Some aims are shared between these two applications: growth in animal-free conditions, fast and high-density growth in single-cell suspension, cell immortalization, and scalability. However, this study also reports preliminary nutritional profiles of the cells to address the specific applications.

### 4.1 MsNAC culture yields immortalized cells that are adaptable to animal-component free media

Cell immortalization (i.e., culturing of cells past their natural point of senescence) is important to ensure sufficient biomass can be produced and cells remain stable throughout the production process. Immortalization is typically achieved either through genetic manipulation or spontaneous immortalization through random mutations over many subcultures. While both methods have been used to generate cells for CM (Letcher et al., 2022; Saad et al., 2022; Stout et al., 2023a), spontaneous immortalization may be favorable from a consumer acceptance and regulatory standpoint (Bryant et al., 2020). MsNACs were spontaneously immortalized and showed stable growth rates past 100 doublings, and a continuous cell culture has been maintained for over 1.5 years after isolation. Cultivation in animal-free conditions is necessary to align with the goals of CM to produce animal-derived products without the animal. Despite many advances in generating animal component-free media for mammalian (Stout et al., 2022) and avian (Pasitka et al., 2023) cells, animal-free media for CM remains an active area of research. MsNACs were easily adapted to commercial animal-component free media (SF900 III) over four passages and all further studies were performed in Sf900 III media.

### 4.2 MsNACs proliferate in high-density suspension culture

After a prolonged selection period for non-adherent cells, MsNACs were transferred to single-cell suspension growth in shake flasks. Single-cell suspension is preferable over adherent growth (either in static culture in flasks or microcarrier methods in dynamic culture) because cells can typically reach higher densities as they are no longer constrained by vessel or carrier surface area (Rubio et al., 2019b). High-density suspension culture without microcarriers is often inhibited by the formation of large aggregates. Although biofilms and some cell aggregations were observed at the endpoint of culture in shake flasks, this did not seem to significantly impact cell growth. This is likely due to the long selection period for anchorage-independent cells. Passaging of cells in suspension was simply performed by diluting the culture, without removing spent media or breaking up cellular aggregates with enzymatic methods as may be necessary with other single-cell suspension lines used for cultivated meat (Pasitka et al., 2023).

One important metric for optimization in CM is the highest achievable cell density. Increasing the cell density (as measured in cells per mL of media) results in less media and electricity usage per unit of cell biomass, thus reducing costs (Humbird, 2021). MsNACs reached an average of ∼27E6 viable cells/mL at passage 35 and ∼20E6 viable (∼30E6 total) cells/mL at passage 50 after cryopreservation without further media or growth optimization. Other insect cells also reach similar densities in single-cell and animal-component free formulations. For example, Sf9 cells reached 14E6 cells/mL in 100-ml Erlenmeyer shake flasks (Käßer et al., 2022), 16E6 cells/mL in 50-mL TubeSpin bioreactors (Xie et al., 2011), and 12-14E6 cells/mL in 1-5 liter shake flasks (Cronin, 2020). Sf9 cells in a 14-L airlift bioreactor reached 10E6 cells/mL in an earlier formulation of Sf900 III media (King et al., 1992). Thus, MsNACs show slightly higher achievable cell densities than high-density insect cell cultures previously reported, demonstrating consistency of insect cells from different species and starting tissues/developmental timepoints to reach high density single-cell suspension in animal-free media.

Further optimization should lead to even higher achievable cell densities. For example, Sf9 cells (the result of a clonal isolation, or separation of heterogeneous cell populations into single cells and propagation of genetically identical clonal populations) can reach approximately double the cell densities of their parent cell line, Sf21 (Cronin, 2020). It is also important to note that while MsNAC cell doubling time during log phase is comparable to other insect cell lines as well as to vertebrate cell lines cultured in suspension, the values were slower than optimized cell lines such as High-Five or Sf9 with doubling times as low as 18 hours (Käßer et al., 2022). As density increased, cell growth rates slowed considerably. Because we observed glutamate depletion and ammonia buildup at densities above 3-4M/mL without media exchange, the slow growth of our high-density cultures was likely due to waste accumulation, nutrient gradients, and altered metabolism. If MsNACs were to be considered for larger-scale culture, growth rates (especially at high density) would need to be optimized, potentially via clonal isolations, media optimization, or genetic engineering.

To our knowledge, the cell type with the highest published cell density in single-cell suspension studied for CM is an embryonic chicken fibroblast cell line, which reached 7E6 viable cells/mL in fed-batch culture with glucose supplemented daily (Pasitka et al., 2023). This yield increased substantially by using a continuous (perfusion) culture system, with cells reaching an impressive 108E6 cells/mL after 15 days (Pasitka et al., 2023). While perfusion was not explored in the present study, we expect that optimization of feeding strategies could enhance yield. Thus, MsNAC growth without optimization was comparable to spontaneously immortalized chicken fibroblasts similar to those used in USDA and FDA-approved CM products. One major difference between MsNAC and traditional CM products, however, are the environmental conditions for growth. Where avian or mammalian cells require 37-29°C incubation temperatures with CO_2 s_upplementation, MsNACs were grown at 27°C without CO_2 (_Pasitka et al., 2023). These growth conditions are typical for insect cells, which have been routinely grown in adjacent industries in 50-liter wave bioreactors, 14 and 21-liter airlift bioreactors, or even retrofitted 150-liter microbial fermenters (Garnier et al., 1996; Ghasemi et al., 2019; Kaiser et al., 2022; King et al., 1992; Maiorella et al., 1988). Biotechnology companies likely scale over 1000-liter bioreactors, although results are not published in peer-reviewed journals (Drugmand, 2007). This precedent positions insect cells (such as MsNACs) well for large-scale production for CM.

### 4.3 Spent media offers insights into cell metabolism

Spent media analysis of MsNACs in batch culture was used to gain insights into metabolism and potential avenues for further optimization. Glucose was assumed to be the primary energy source for insect cells via glycolysis and cellular respiration (Drugmand et al., 2012). In animal cell culture, the consumption of glucose is typically linked with the production of lactate as a waste product during anaerobic metabolism. Interestingly, during 7 days of culture (with growth slowing at around day 5), glucose was not consumed, and lactate was minimally produced, indicating that the cells were not relying on glycolysis as a primary energy source despite the relatively high levels of glucose (∼9 grams/liter) present in the media. This contrasts with typical insect cell metabolism. Although Sf9 cells *can* consume non-glucose carbohydrates such as maltose or fructose (Käßer et al., 2022), if glucose is present both Sf9 and High-Five cells exhaust it as a carbon source (Guardalini et al., 2023; Rhiel et al., 1997). Insect cells produce significantly lower levels of lactate in comparison with mammalian cells (Guardalini et al., 2023), although High-Five cells have been shown to produce lactate up to 16 mM in suspension. It is hypothesized that this accumulation is due to aggregate formation (and therefore low-oxygen environments for cells) as High-Five cells grown in single-cell suspension do not accumulate this level of lactate (Ikonomou et al., 2003; Ikononou et al., 2001; Yang et al., 1996). Thus, MsNACs show an altered metabolic pathway for energy production compared with commonly used insect cell lines and further exploration into this metabolic phenomenon is warranted.

In addition to glucose and lactate, the amino acids glutamine and glutamate, along with their metabolic byproduct ammonia were assessed. Although insect (and mammalian) cells are able to biosynthesize glutamine, making it a nonessential amino acid, glutamine is considered essential in many cell culture systems because glutamine consumption rate exceeds the biosynthesis rate (Mitsuhashi, 1982; Yoo et al., 2020). Glutamine plays a vital role as the primary nitrogen donor for amino acid and nucleotide syntheses, undergoing conversion to glutamate and subsequently to alpha-ketoglutarate before entering the TCA cycle. The deamination of glutamine to glutamate and further to alpha-ketoglutarate results in the release of ammonia. The buildup of ammonia (alongside lactate) is toxic to cells, necessitating media refreshment to sustain continual cell growth (Hubalek et al., 2023).

Metabolic waste buildup is one of the biggest limiting factors for large-scale biopharmaceutical production – with 2-10 mM ammonia and ∼20-100 mM lactate inhibitory levels for mammalian cells (Humbird, 2021). Models of Chinese Hamster Ovary cells using these inhibitory levels and the production levels with metabolically optimized cells is still hypothesized to lead to culture systems that prohibit an economically sufficient cell density for CM (Humbird, 2021). Surprisingly, MsNACs consumed minimal glutamine over 7 days, but still produced significant amounts (up to 15 mM) of ammonia. In contrast, glutamate was nearly exhausted from the medium over 7 days and was likely the source of the ammonia production. Again, nutrient consumption patterns of MsNACs appear to differ from commonly used insect cell lines such as Sf9 cells, which are known to consume high levels of glutamine (Bédard et al., 1993; Benslimane et al., 2005). For example, Sf9 cells cultured in SF900 III media had the highest specific consumption rate for glutamine of nutrients studied in stirred tank culture and produced a maximum of 6.1 mM ammonia – nearly half the amount produced by MsNACs (Guardalini et al., 2023). Because MsNACs were negatively affected by ammonia concentrations above 6.25 mM, we hypothesize that the rapid accumulation of ammonia is at least partially responsible for the cessation of growth.

In an effort to reduce ammonia production and further understand MsNAC metabolism, preliminary experiments were performed in SF900 III media with reduced amounts of glutamine. When glutamine was reduced to ∼0.3 mM, the cells produced less ammonia and reached similar maximum VCDs compared with cells grown in control media (which contains ∼14 mM glutamine) (SI Figure 3F,G). While further optimization is necessary to understand how to further increase batch-culture VCD, the observation that MsNACs can proliferate with significantly reduced glutamine is favorable in the context of CM. An entomoculture-based TEA showed that in multiple media formulations L-glutamine is an order of magnitude more costly than glutamate (Ashizawa et al., 2022). Glutamine is also known to easily break down when stored for long periods, releasing inhibitory ammonia into the media. While a stabilized version of L-glutamine is used in many mammalian culture systems (l-alanyl-l-glutamine), insect cells have difficulty metabolizing l-alanyl-l-glutamine, therefore L-glutamine is still the preferred substrate in commercial media (ThermoFisher, personal communication). Because of this breakdown of glutamine, plant hydrolysates are hypothesized to be incomplete amino acid sources (Humbird, 2021). A 2021 TEA suggested that soybean meal could be a source of all essential amino acids for mammalian cell culture, except for glutamine and tyrosine. However, this meal produced more than double the required glutamate, and thus could be explored as a potential media supplement for MsNACs (Humbird, 2021). MsNACs were also able to proliferate and reach similar VCD to control media with *both* glutamine and glucose reduced media (0.1 g/L and 0.5 mM, respectively) (SI Figure 3F). These results show the adaptability of the cells to growth on different substrates, representing the potential for further media optimization that should be explored in future work.

Ingredients for CM are assessed not only for their monetary cost, but also their environmental cost to produce and their availability to access at scale. Unfortunately, due to the proprietary nature of the cell culture media in this study we cannot determine the environmental cost and availability of all components in the media. It is important to note that commercial insect media formulations such as the SF900 III media used in this study are designed for insect cells that are producing recombinant proteins. The increased metabolic burden of protein synthesis leads to increased energy consumption, meaning the media likely contains an excess of nutrients than needed for cell growth vs. optimized recombinant protein production (Käßer et al., 2022). Future studies should explore customized media for MsNACs using only the nutrients necessary for growth to increase efficiency and decrease cost – for example, using design-of-experiments to create optimized formulations (Saisud et al., 2023). The ideal media formulation would primarily rely on plant or fungal hydrolysates instead of individually produced amino acids, as the high cost of amino acids are prohibitory to large-scale production (Humbird, 2021). For example, rapeseed proteins and peptides (a byproduct of canola oil production) have also been shown to increase Sf9 cell density by 60% in serum-free conditions (Deparis et al., 2003) and have been explored in mammalian cells as a low-cost serum-free media supplement (Stout et al., 2023b).

### 4.4 Nutrition

#### 4.4.1. Proximate analysis

In addition to growth kinetics, adaptability to different culture vessels, and spent media analysis, a basic nutritional profile was compiled for MsNACs including proximate analysis as well as amino acid, mineral, and fatty acid composition. One of the advantages of the ability to easily scale to a 2.4-liter bioreactor was the feasibility of generating sufficient biomass (15-20 grams of wet cell weight per sample) for food composition assays. These samples were used for proximate analysis to determine the proportion of protein, fat, carbohydrates, and ash in each batch of cells. These analyses showed that the dry weight nutritional breakdown of MsNACs was similar to FDA and USDA-approved embryonic chicken fibroblasts. Results can also be put in context of other alternative protein sources: whole-insect powders or “flours” are gaining popularity, with the most common coming from mealworms or crickets. Dry weight proximate analysis shows that protein content from these powders are lower than MsNACs, with 65.5 +/- 0.5% and 66 +/- 0.3% for cricket and mealworm, respectively (Stone et al., 2019). The higher protein content in MsNAC dry weight compared with insect powders may be because of the lack of chitinous exoskeleton from whole insects, indicating that pure embryonic cell biomass may be more efficient for protein production than whole insects. Plant-based protein sources from fava beans and yellow pea concentrates have even lower protein content, at 62.5 +/- 0.6% and 55.1 +/- 1.2%, respectively (Stone et al., 2019). Future studies should include analysis of the in vitro protein digestibility to ensure that the protein is bioavailable (Stone et al., 2019).

#### 4.4.2. Amino acids

Beyond proximate analysis, separate samples were analyzed for amino acid breakdown compared with DF-1 (embryonic chicken fibroblast) cells. While the overall breakdown was similar, DF-1 cells cultured in-house had increased molar concentrations of lysine, threonine, and valine. One 20-gram wet cell weight sample was also analyzed via acid hydrolysis and compared with cultivated embryonic chicken fibroblasts from two companies as well as their published values of conventional chicken. Again, overall amino acid breakdown was similar when analyzed as percentage of total amino acids reported. Unfortunately, lysine was not analyzed with the acid hydrolysis method and further analysis on the high lysine content of DF-1s compared with MsNACs found via LC-MS/MS should be pursued.

#### 4.4.3. Minerals

Mineral content was also analyzed, as meat is an important source of essential minerals which are necessary nutrients to support cell function and growth (Falowo, 2021). Minerals can also contribute to the taste, texture, and appearance of meat products (Falowo, 2021). Interestingly, MsNACs had significantly more iron and calcium and increased, but not significant, levels of zinc, copper, magnesium, and phosphorus. These data are consistent with whole-animal nutritional analysis that shows insects are a good source of these minerals (Finke, 2002; Montowska et al., 2019; Stone et al., 2019). Just as mineral composition of whole animals largely depends on diet, cell-based mineral content will be influenced by media composition as well as cells’ ability to uptake each mineral. The animal-free SF900 III media therefore showed favorable mineral composition, with MsNACs able to uptake each mineral analyzed with equal or higher concentrations compared with DF-1 cells. This data is consistent with previous research using a *Drosophila* (fruit fly) cell line, which showed that when adjusting for cell size, insect cells had more iron and zinc compared with a mammalian (mouse myoblast) cell line. This research also found that using iron-fortified serum increased iron density in *Drosophila* cells (Rubio et al., 2019a). While serum should be avoided, similar mineral fortification could be pursued if one wanted to nutritionally enhance MsNACs, for example with iron-fortified yeast extract (Sabatier et al., 2017). While initial results are promising, further analysis on the bioavailability of MsNAC minerals compared with whole insects (or other meat sources) is warranted, as the exact source of the iron is unclear because the SF900 III formulation is proprietary. Although many minerals are essential to a healthy diet, accumulation of heavy metals are a growing concern for consumption of whole insects for food and feed, where insects sequester toxic heavy metals from feeding on agricultural waste (Malematja et al., 2023). While exposure to low levels of heavy metals such as lead, mercury, chromium, cadmium, and arsenic are inevitable, food must be closely monitored to avoid metal poisoning (Balali-Mood et al., 2021). The high level of control in cell culture can decrease the risk of heavy metal accumulation – unsurprisingly, there were no detectable levels of the heavy metals included in our analyses of MsNACs (lead, arsenic, and cadmium).

#### 4.4.4. Fatty acids

Finally, fatty acid content was analyzed and compared between DF-1 and MsNACs. Fatty acid data here is likely less relevant for both DF-1 and MsNACs, which are undifferentiated cells intended to be the animal protein in a product that would likely be supplemented with either plant-based fats or a different lipid-accumulating cell. Although not explored here, embryonic *M. sexta* cells have been investigated as a cultured fat source using a soybean oil emulsion to induce lipid accumulation – it is likely that MsNACs would be able to uptake lipids in a similar manner and could act as a fat source (Letcher et al., 2022). Although the data showed high variability, the majority of free fatty acids were not significantly different between the two cell types. The MsNACs generally had higher lysophosphatidylcholine and the DF-1s had higher phosphatidylcholines, representing differences in the insect and chicken cell membrane composition. Because phospholipids are responsible for the majority of meat-specific flavor, further investigation into potential differences in flavor between MsNAC and DF-1 (and other CM or meat sources) is warranted (Huang et al., 2010; Igene and Pearson, 1979).

#### 4.4.5. Nutrition caveats

Although comparisons between DF-1 cells and MsNACs were intended to put nutritional data in context, it is important to note that comparisons between cell types are difficult due to cultivation in different media (DF-1 cells were cultured in media containing 10% FBS) which may impact the nutritional profile. Overall, it is difficult to determine how MsNACs may compare to other CM cell types based on limited and conflicting nutritional data and variability in culture conditions. Further, MsNACs are not intended to be a “complete” product, rather a potential ingredient and proof-of-concept that insect cells have favorable baseline nutritional properties that are on-par with currently the only FDA and USDA-approved cell type. We envision that MsNACs or similar insect cell lines may be valuable to the CM industry combined with plant-based products to provide animal nutrition and taste for human consumption.

## 5. Limitations & Future Work

Many additional improvements could be pursued to further characterize and engineer MsNACs. This could include altering media formulations, genetic engineering, or clonal isolations to select for even faster-growing cells that reach higher densities. Growth in bioreactors larger than 10 liters should also be pursued to ensure cells can be scaled effectively. Nutrition could be enhanced by media supplementation or use of the baculovirus expression system to increase protein content or express specific nutrients (Verkerk et al., 2007). Further studies are also required to assess the bioavailability of MsNAC nutrients compared with traditional meat. Larger scale culture also allows for more robust flavor/sensory analysis, which could include analysis of volatile compounds through GC-MS or sensory panels with human participants.

Beyond further engineering and characterization of MsNACs, it is important to consider potential biases in the cell line development process. Here, we described selection for three main traits that are favorable for CM during the expansion of MsNACs: 1) cells that grow well in animal component-free media, 2) cells that grow in single-cell suspension, and 3) cells that naturally spontaneously immortalize. Each selective pressure likely led to cultivation of a subpopulation of cells that is not representative of the originally isolated population. Although these selective pressures led to a robust cell line with stable growth over 1.5 years, it is possible that other desirable traits were lost in the process.

Starting from an embryonic cell population provides significant flexibility in selection criteria because many different early-stage cell types are present. Other selection criteria could include cells that grow in higher densities, in simpler media formulations, at lower temperatures, or cells that are more metabolically favorable (e.g., produce less ammonia or are less sensitive to ammonia). Future studies could even use clonal isolations to determine the most nutritious or palatable cell line.

Finally, consumer acceptance studies will need to be performed to determine the attitude of consumers towards cultured insect protein incorporation into their diet. Previous work has shown that Western consumers rate dishes that contain visible insects poorly (e.g., “fried mealworm” or “locust salad”), while disguised insect material (e.g., “pizza containing protein derived from insects”) were rated higher (Schösler et al., 2012). Thus, with appropriate marketing, insect CM could bypass Western consumers’ food neophobia associated with entomophagy. If consumer acceptance proves too difficult, insect cells could also be incorporated into pet food, which accounts for approximately 20% of global meat consumption (Knight, 2023).

## 6. Conclusions

The present study provides a strategy to generate insect cells that are spontaneously immortalized, grow in animal-component free media, and proliferate in high densities in single-cell suspension culture in multiple different culture vessels; all of which remain active challenges in scaling up CM production. The nutritional composition of the cells was also investigated and was comparable to embryonic chicken fibroblasts. While consumer acceptance of insect cell-based cultured meat may be a challenge in Western communities, this study offers insight into how insect cells may be incorporated into large-scale cultivated meat production processes with potentially lower production costs yet comparable nutritional benefit.

## Abbreviations

MsNAC: *Manduca sexta* non-adherent cells;
CM: Cultivated Meat;
VCD: Viable Cell Density;
FBS: Fetal Bovine Serum

## Data availability

The authors declare that data supporting this study are available within Supplementary information. Extra data are available from the corresponding author upon request.

## Author Contributions

**Sophia M. Letcher**: Conceptualization, Investigation, Methodology, Visualization, Formal Analysis, Writing – Original draft, Validation. **Olivia P. Calkins:** Methodology, Validation, Formal Analysis, Writing – Review & Editing. **Halla Clausi:** Validation, Formal Analysis, Writing – Review & Editing. **Aidan McCreary:** Validation, Formal Analysis, Writing – Review & Editing. **Barry Trimmer:** Conceptualization, Supervision, Writing – Review & Editing. **David L. Kaplan:** Conceptualization, Funding acquisition, Supervision, Writing – Review & Editing.

## Acknowledgements

We thank New Harvest for their support of this work. We also thank the Metabolomics Innovation Center and Eurofins for assistance with nutritional testing, Chemometec for assistance with cell counting, and Ark Biotech for the support with bioreactor runs. Thank you to ThermoFisher for their generous donation of customized Sf900 III through their academic partnership program. Thanks also to Ellie Contreras for the donation of DF-1 cells, Naya McCartney for assistance with collecting *Manduca sexta* eggs for isolations, and Natalie Rubio for insect cell culture guidance. Thanks to Yu-Ting Dingle for education on Adobe Illustrator for figure preparation. Finally thank you to Tufts University Center for Cellular Agriculture past and present members Michael Saad, Adham Ali, Olympe Jean, and others for scientific input and support.

## Supplemental Information

### Supplemental Materials & Methods

#### Species Validation

Genomic DNA was extracted from MsNACs from three different passages using GeneJET Genomic DNA Purification Kit (ThermoFisher Scientific #K0721) following manufacturer’s instructions. Primers were designed to target *Manduca sexta* cytochrome c oxidase subunit I (Forward: 5’-GGAGCTGGTACAGGTTGAACAG – 3’, Reverse: 5’-TGGATCTCCCCCTCCAGCAG-3’) and PCR was performed using Q5^®^ High-Fidelity 2X Master Mix (New England Biolabs (NEB) #M0492, Ipswich, MA, USA). Successful PCR was confirmed by running the PCR product on a 2% agarose gel and imaging using the G:BOX Chemi XR5 (Syngene, Bangladore, India) and a band of the expected product size was identified. PCR products were then purified using Monarch^®^ PCR & DNA Cleanup Kit (5 μg) (NEB, # T1030), and the forward primer was used for amplification using Sanger sequencing (GENEWIZ from Azenta, South Plainfield, NJ). The sequences were aligned to known COI sequences for *Manduca sexta,* as well as other related lepidoptera species (*Bombyx mori, Spodoptera frugiperda,* and *Manduca quinquemaculata)* using NCBI Nucleotide BLAST.

#### Mycoplasma Test

MycoAlert assay (LT07, Lonza) was used to detect potential mycoplasma activity. Cell culture was centrifuged at 380 *xg* for 5 minutes and supernatant was transferred into a 96-well plate. Samples were incubated alongside positive and negative controls with reagent solution for 5 minutes and luminescence was measured on a microplate reader.

Samples were then incubated with a substrate solution for 10 minutes and luminescence was measured again. The second reading was divided by the first reading to generate the luminescence ratio. Samples were measured in duplicate and controls in triplicate. Samples were collected from passages 62, 46, 32, and 33. Negative control was fresh Sf900 III media.

#### 10-liter Bioreactor Run

The 10-liter airlift bioreactor was operated similarly to bioreactor runs described in the main text. The 2.4-L bioreactor was used to inoculate the 10-L bioreactor at a viable cell density of 500,000 cells/mL. Culture media was supplemented with 0.5% Antibiotic-Antimycotic (15240096, ThermoFisher Scientific). The bioreactor was operated at 27C with an air sparge rate of 0.4 LPM. 1-2mL of a 1:100 mixture of an antifoam agent and RODI was added as needed to minimize foaming levels. When ammonia levels reached 10 mmol/L, 50% of the culture media was exchanged. Half of the total working volume of media was removed and spun down at 325xg for 13 minutes at an acceleration level of 9 and deacceleration level of 6. The resulting cell pellet was then resuspended in fresh media before being reintroduced into the bioreactor.

#### Fed Batch Culture

MsNACs were seeded at 500,000 cells/mL in 30 mL working volumes of Sf900 III media in 125-mL vented shake flasks (ThermoFisher <u>4113-0125</u>) in two sets of triplicates. After cells had reached approximately 3M cells/mL, one set of flasks had a 50% media exchange every other day (“Media Exchange”), and the other set of flasks had 5 mL of Sf900 III media added every other day (“Fed-Batch”). Total cell counts and viable cell density were taken daily using an automated cell counter (NC-200™, Chemometec).

#### ExpiSF vs. SF900 III Growth Curves

MsNACs were seeded at 500,000 cells/mL in 25-50 mL working volume of either SF900 III or ExpiSF CDM (ThermoFisher A3767801) in 125-mL shake flasks (ThermoFisher <u>4113-0125</u>) in triplicate. Total cell counts and viable cell density were taken daily using an automated cell counter (NC-200™, Chemometec).

#### Nutrient Deprivation Studies

For studies involving nutrient removal (either L-glutamine and/or glucose), Sf900 III was custom-made by ThermoFisher. For all studies involving custom media, the media used for control conditions were Sf900 III minus L-glutamine + 13.7 mmol/L of sterile-filtered L-glutamine () to ensure a controlled starting concentration. Similarly, media conditions without glucose were made by adding 13.7 mmol/L of sterile-filtered L-glutamine to the Sf900 III minus L-glutamine, minus glucose media. Cell-free media controls were included to determine the rate of nutrient breakdown or accumulation due to low levels of media evaporation and were subtracted from final calculations. Growth curves were performed in batch with cells seeded at 500,000 cells/mL in 30 mL working volumes, and cell count/viability and spent media samples were collected at day 0, 3, 4, 5, and 6 to capture log phase growth. Spent media samples were stored at -80C until analysis as described in main text.

**SI Figure 1.**
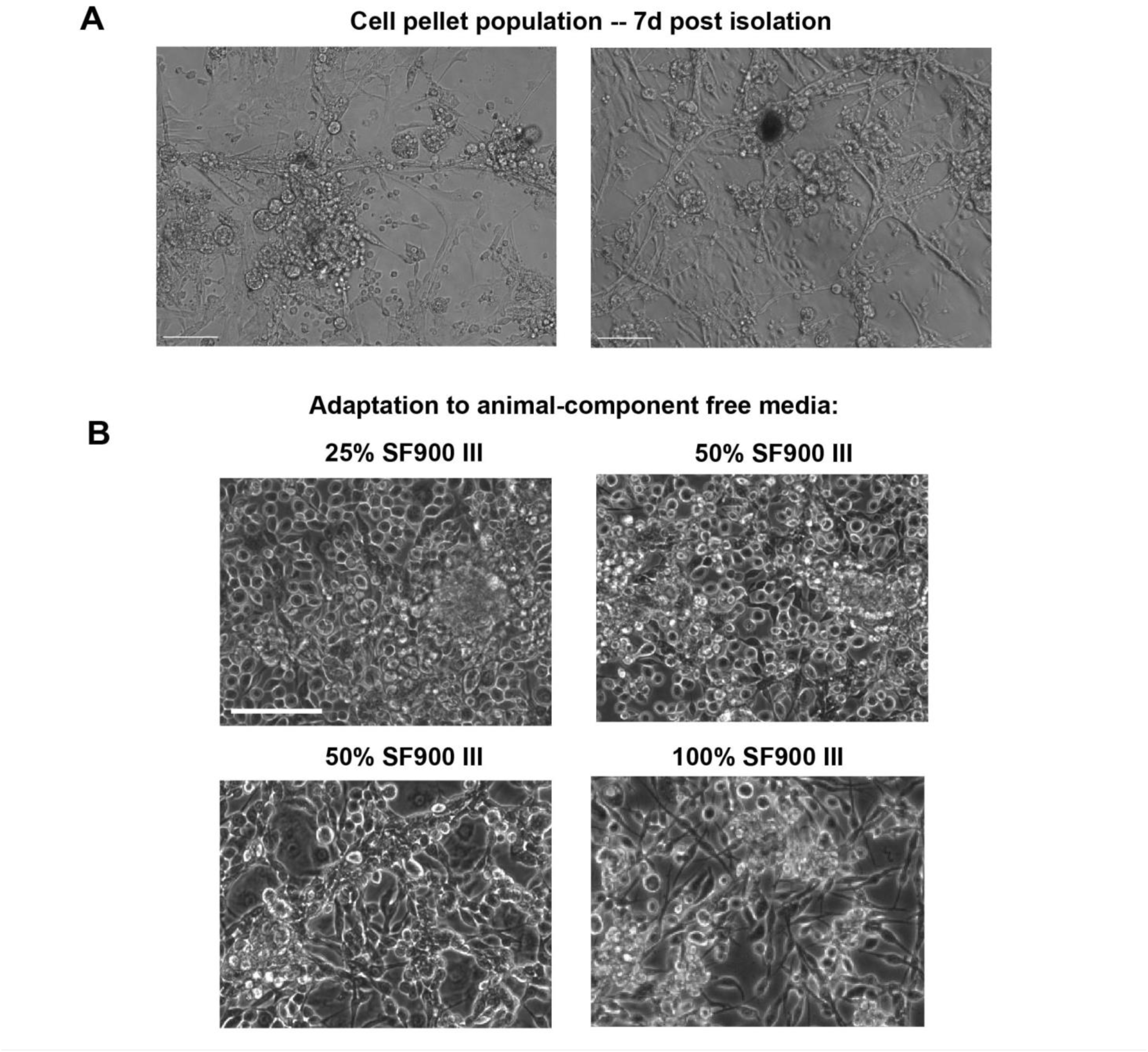
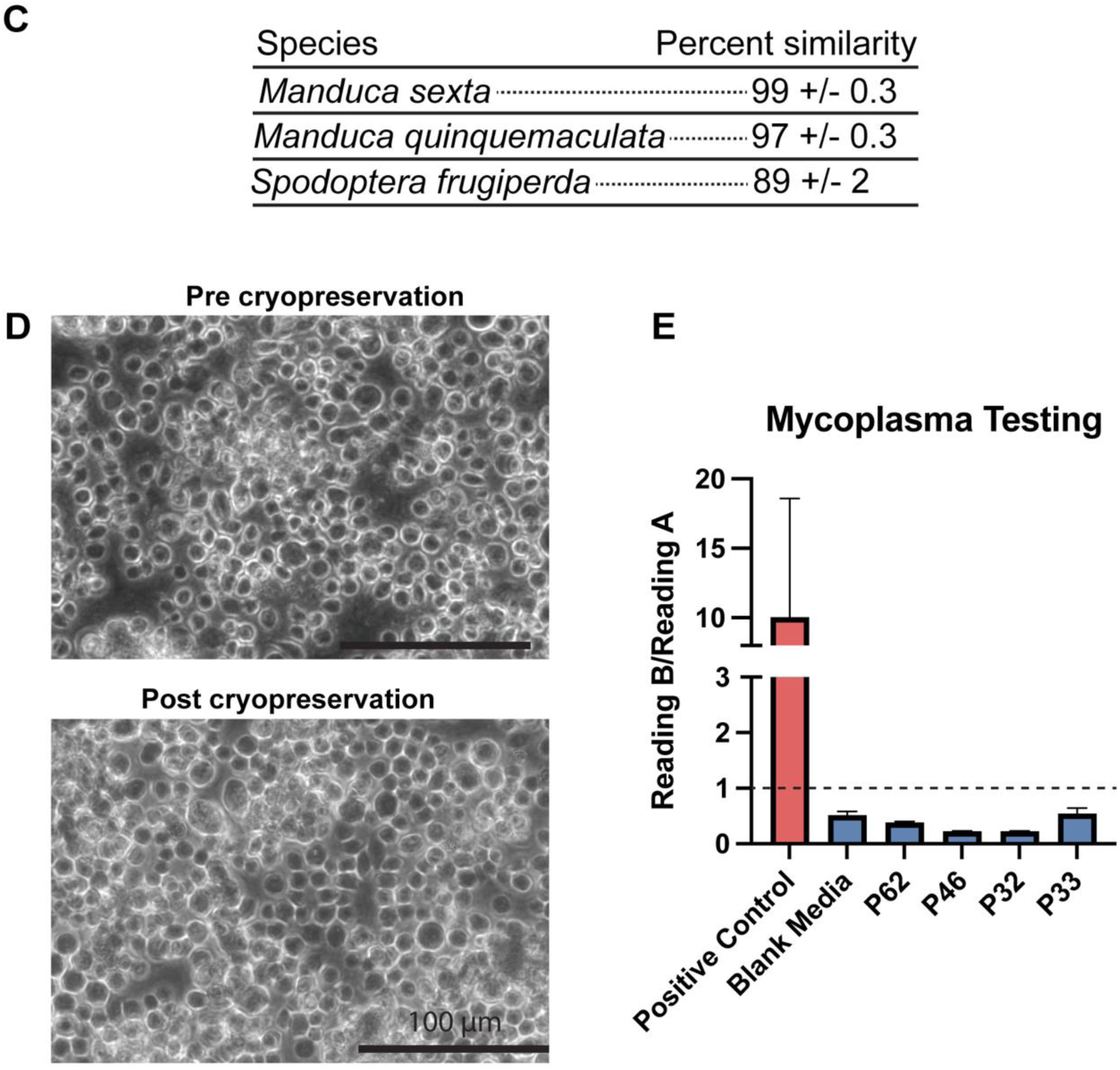
(A) Phase contrast images of two fields of view of the cell pellet population, showing a range of cell morphologies. (B) MsNAC morphology while being adapted to SF900 III media from media containing 10% FBS over four passages, each with increasing amounts of SF900 III media. SB = 100 microns. (C) Species validation via sanger sequencing compared to published COI genes for MsNACs compared with *Manduca sexta, Manduca quinquemaculata,* and *Spodoptera frugiperda.* (D) phase contrast images before (top) and after (bottom) cryopreservation in SF900 III media SB = 100 microns. (E) Mycoplasma testing of passage 62, 46, 32, and 33 compared with blank Sf900 III media and positive control. Values above 1 represent the presence of mycoplasma.

**SI Figure 2.**
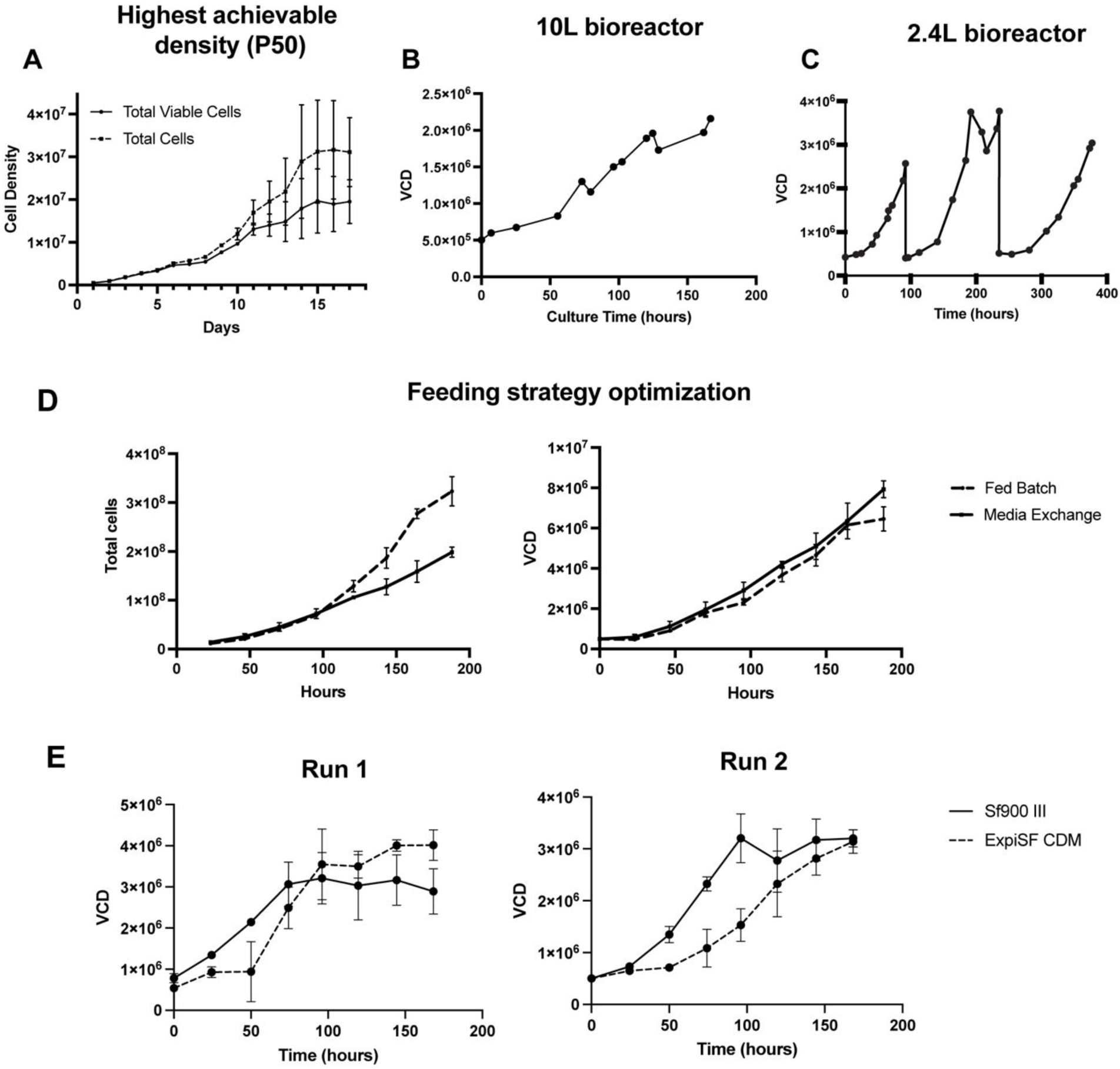
(A) Total viable cells and total cells from passage 50 MsNACs grown over 17 days with 50% media exchange every other day (n=3) (B) MsNAC growth in 10-liter bioreactor over 150 hours. (n=1) (C) Data from Figure 2D with all collected datapoints shown. (D) Comparison between total cells generated from 125 mL shake flasks in either fed batch (adding 5 mL each day without removing media) and 50% media change (removing and replacing 50% of media volume every other day) (left) and viable cell density for each condition (right) (n=3) (E) Growth of MsNACs in 125 mL shake flasks in Sf900 III (solid line) compared with ExpiSF CDM (dashed line). Left run represents cells grown with a 25 mL working volume, while right run represents cells grown in a 50 mL working volume. n=3 flasks per condition.

**SI Figure 3.**
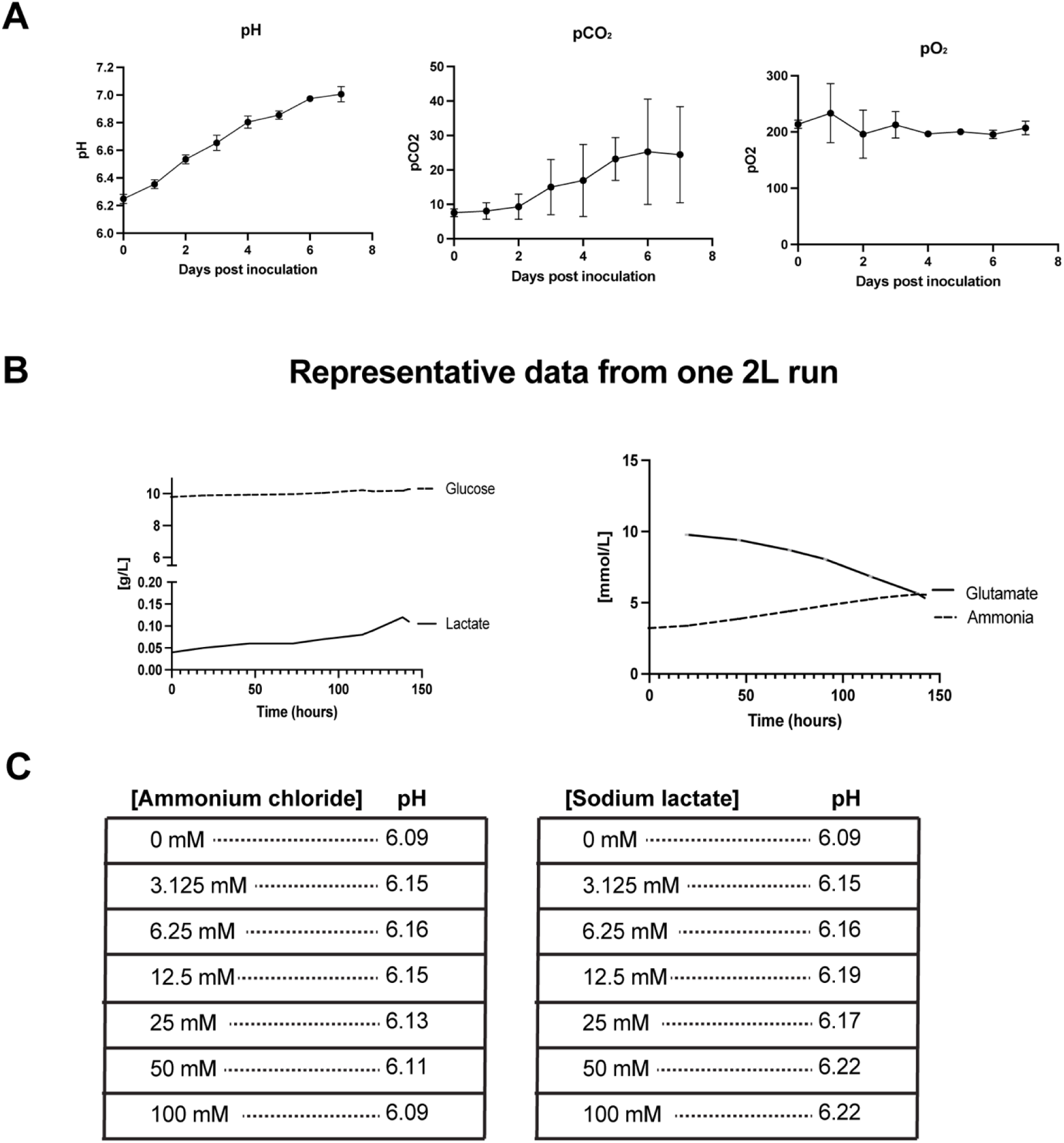
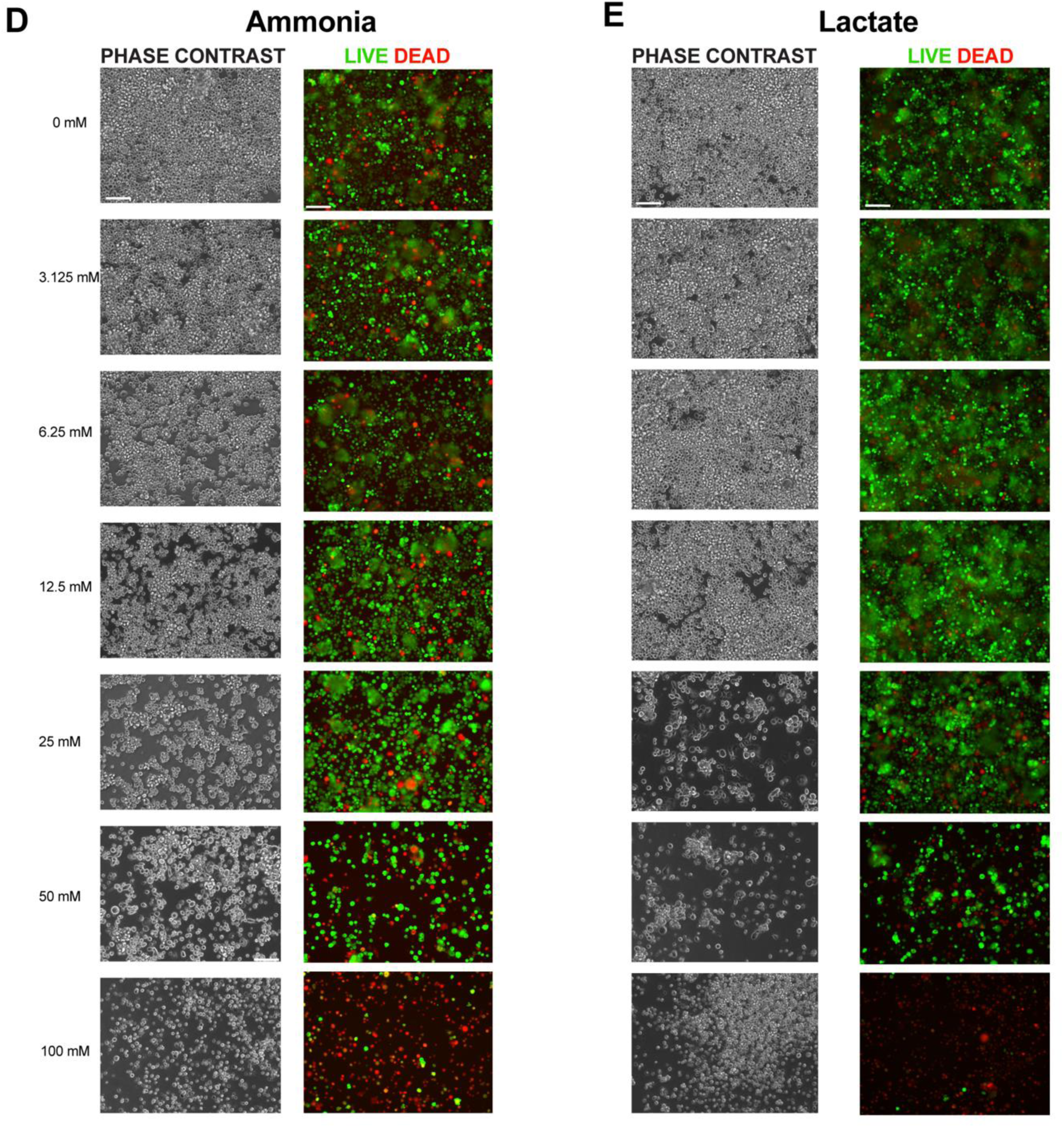
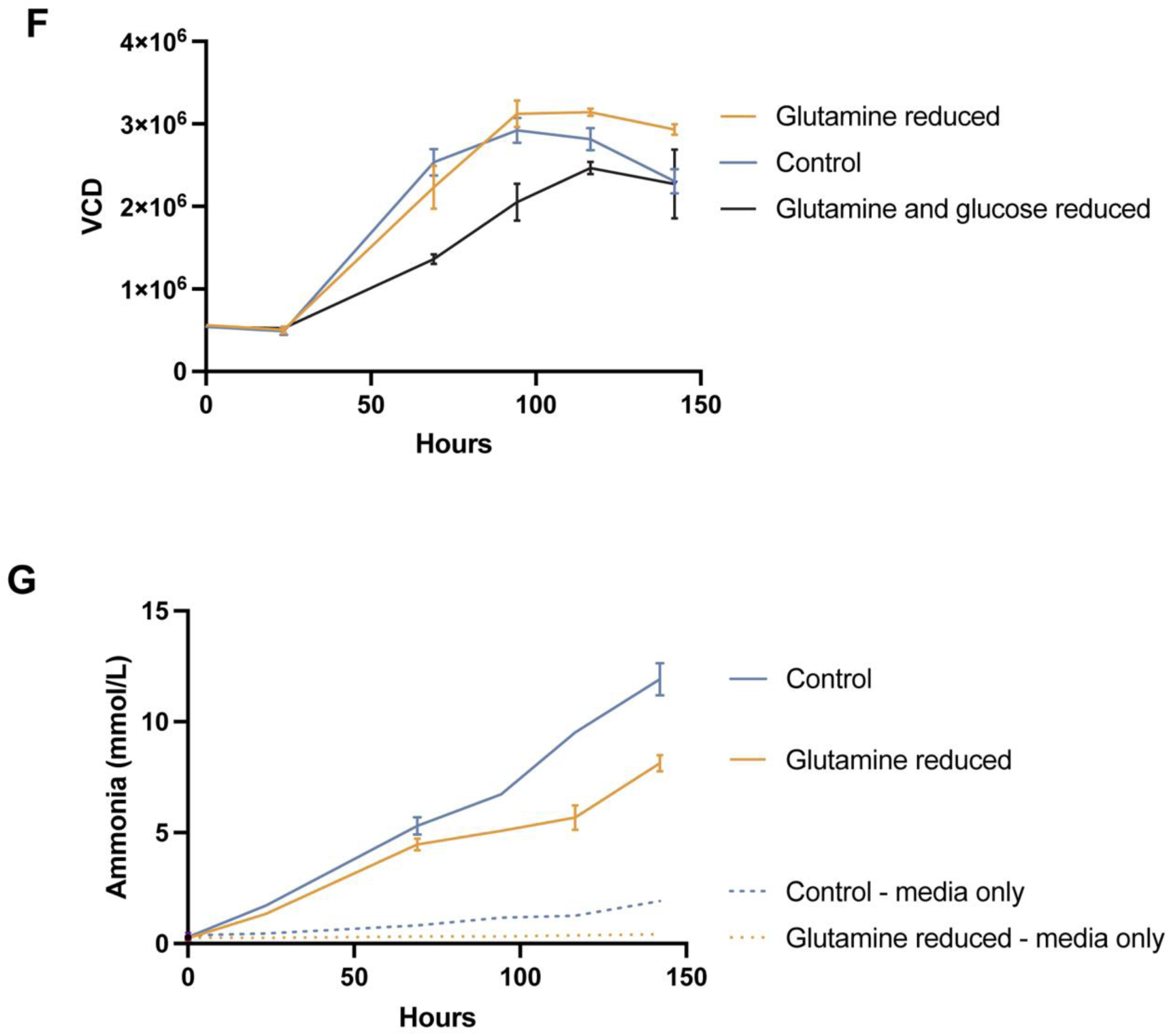
(A) Spent media analysis for pH, dissolved carbon dioxide, and dissolved oxygen for cells grown in Figure 2B (n=3). (B) Spent media analysis for glucose and lactate (left) in g/L and glutamate and ammonia (right) measured in mmol/L over 150 hours of culture (n=1). (C) pH measurements for media used in Figure 3B-E. (D) Phase contrast images (left) and live/dead images (right) of MsNACs after 7 days of culture with increasing amounts of ammonium chloride. (E) Phase contrast images (left) and live/dead images (right) of MsNACs after 7 days of culture with increasing amounts of sodium lactate. (F) Growth over 6 days in glutamine reduced media (∼0.3 mmol/L), control media (containing ∼12 mmol/L glutamine and ∼10g/L glucose) and glutamine and glucose reduced media (containing ∼0.3 mmol/L and ∼0.05 g/L, respectively). *n=3.* (G) Ammonia production in control (blue) and glutamine reduced (yellow) media, with dotted lines showing ammonia accumulation in cell-free media due to amino acid breakdown and/or evaporation.

**SI Figure 4.**
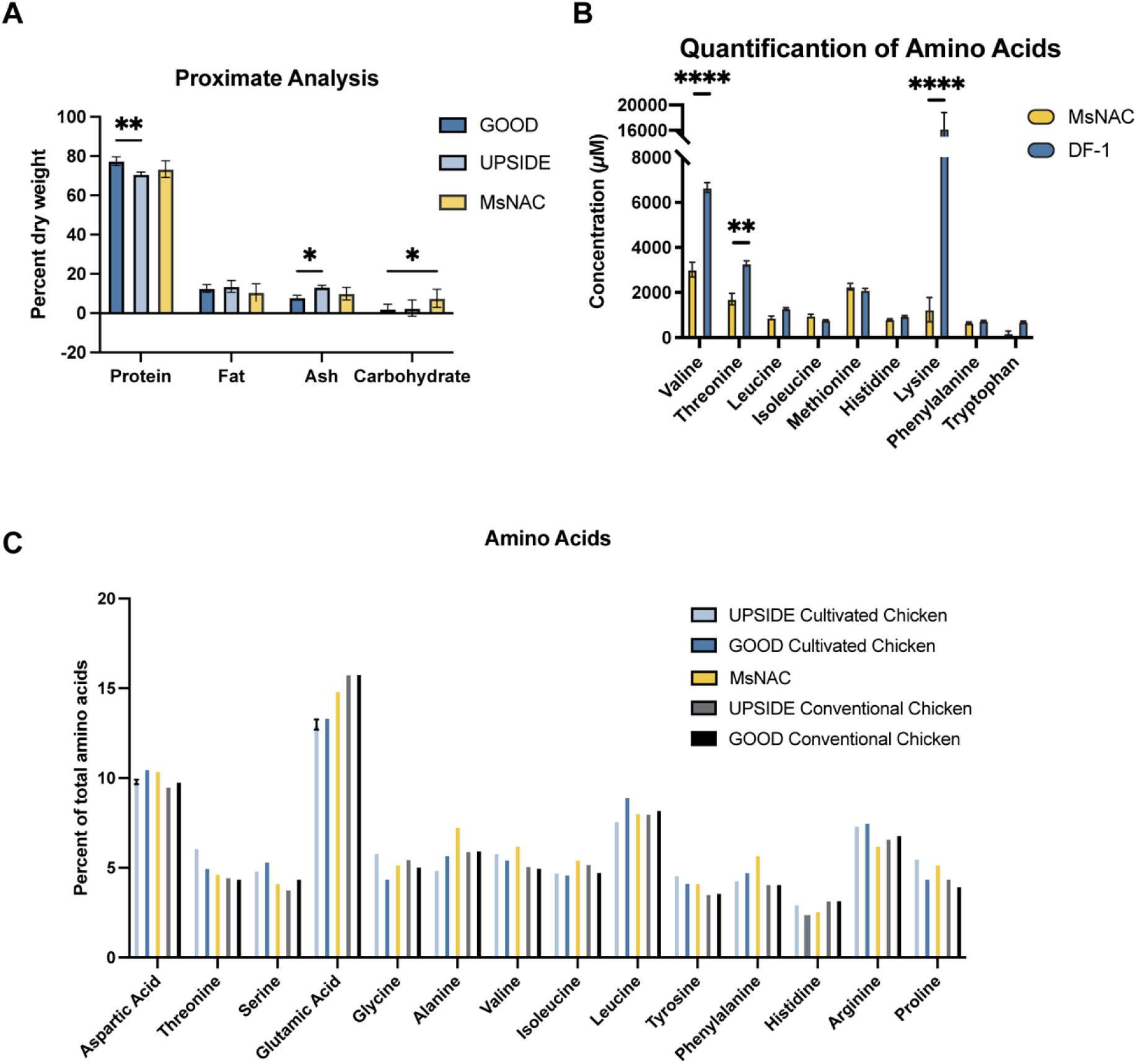
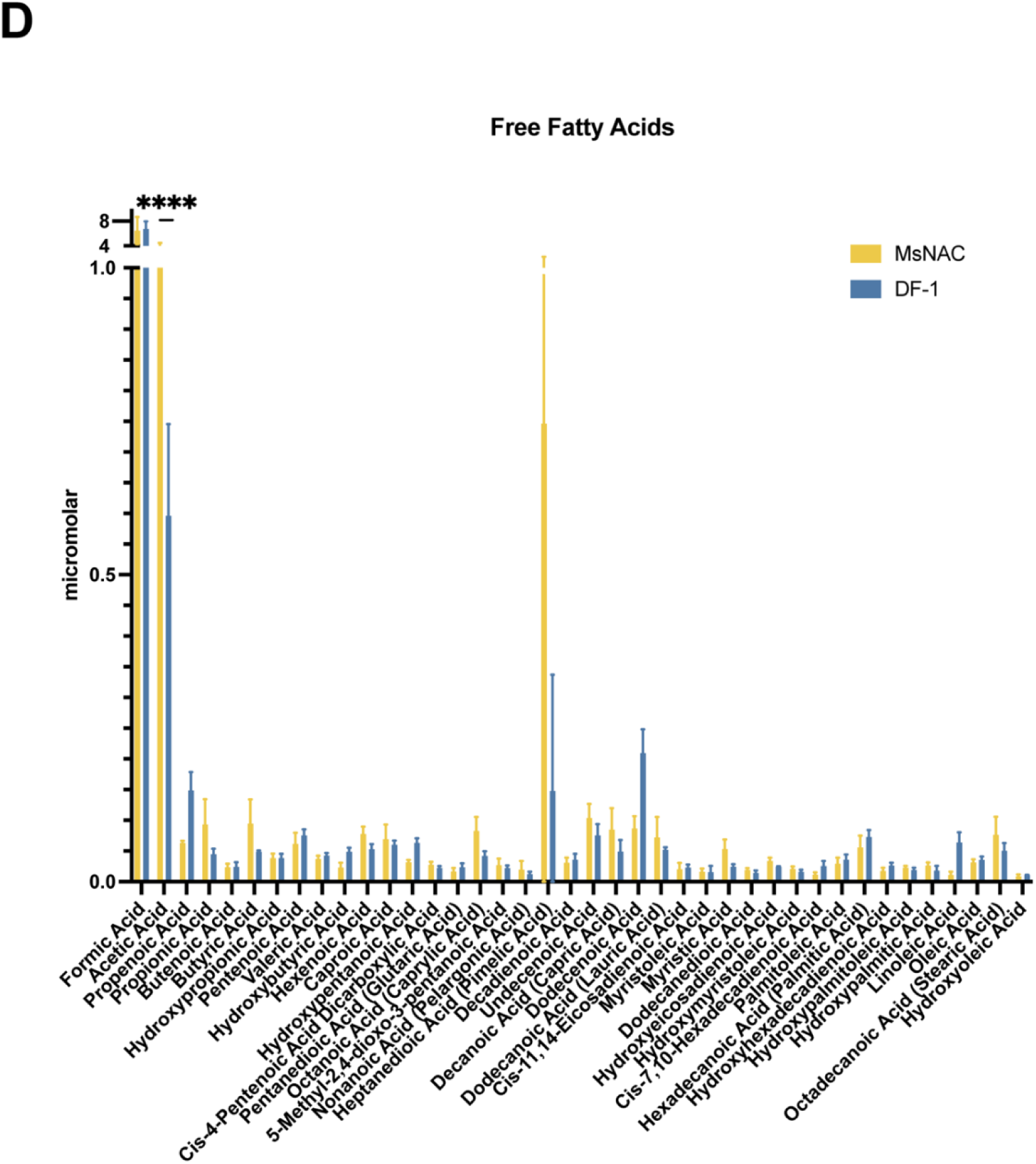
(A) comparison between dry weight protein, fat, ash, and carbohydrate breakdown of MsNAC samples from 5A and GOOD meat’s safety dossier using DF-1 cells and UPSIDE’s dossier using embryonic chicken fibroblasts^1^. (B) micromolar concentrations for essential amino acid breakdown shown in 5E. n=3, 1-way ANOVA with **<0.01, ****<0.0001. (C) comparison between MsNAC (yellow) amino acids analyzed via acid hydrolysis with GOOD and UPSIDE’s safety dossier documenting their product (light and dark blue) and conventional chicken (grey and black). Results are shown as normalized percentages of select amino acids. (D) Comparison of free fatty acid micromolar concentrations between MsNAC (yellow) and DF-1 (blue) 1-way ANOVA with ****<0.0001.

**Supplemental Table 1.** Doubling times over various passages and culture conditions.

| **Passage/culture vessel** | **Average cell doubling time (hours)** | **Doubling time standard deviation (hours)** |
| --- | --- | --- |
| Shake flask - passage 30-35 | 42.64 | 13.85 |
| Shake flask – passage 80-95 | 33.5 | 6.3 |
| 2.4L airlift bioreactor – entire batch | 44.32 | 8.77 |
| 2.4Lirlift bioreactor – log phase | 23.6 | 7.9 |

**Supplemental Table 2.** Numerical values of acid hydrolysis analysis of MsNACs.

| Eurofins AA analysis |  |
| --- | --- |
| Parameter | Result (percent) |
| Alanine | 0.28 |
| Arginine | 0.24 |
| Aspartic Acid | 0.4 |
| Glutamic Acid | 0.57 |
| Glycine | 0.2 |
| Histidine | 0.1 |
| Isoleucine | 0.21 |
| Leucine | 0.31 |
| Phenylalanine | 0.22 |
| Proline | 0.2 |
| Serine | 0.16 |
| Threonine | 0.18 |
| Total Lysine | 0.36 |
| Tyrosine | 0.16 |
| Valine | 0.24 |

**Supplemental Table 3.** Numerical values of essential amino acids analyzed via UPLC-MRM/MR (n=4)

| Essential AA (uM) |  |  |
| --- | --- | --- |
| Parameter | Average | SD |
| Valine | 3017.5 | 385 |
| Threonine | 1703.5 | 196 |
| Leucine | 877.5 | 86.4 |
| Isoleucine | 961 | 71.7 |
| Methionine | 2260 | 163 |
| Histidine | 804 | 28.2 |
| Lysine | 1236.25 | 522 |
| Phenylalanine | 660.25 | 26.4 |
| Tryptophan | 192 | 95.6 |

**Supplemental Table 4.** Numerical values for metals in micromolar for MsNACs analyzed via ICP-MS (n=4)

| Metals (uM) |  |  |
| --- | --- | --- |
| Parameter | Average | SD |
| Sodium | 1.24E+06 | 5.66E+05 |
| Magnesium | 9.78E+03 | 3.71E+03 |
| Phosphorus | 8.40E+04 | 2.77E+04 |
| Potassium | 1.51E+04 | 2.83E+04 |
| Calcium | 2.35E+03 | 9.05E+02 |
| Iron | 1.52E+02 | 5.65E+01 |
| Zinc | 2.31E+02 | 9.78E+01 |
| Chromium | 1.12E+00 | 3.03E-01 |
| Cobalt | 2.94E-01 | 7.08E-02 |
| Copper | 9.91E+00 | 4.89E+00 |
| Selenium | 4.15E+00 | 2.34E+00 |
| Rubidium | 1.15E+01 | 4.86E+00 |
| Strontium | 6.86E-01 | 5.73E-01 |

**Supplemental Tables 5-6.** See attached excel sheet for fatty acid measurement comparisons between DF-1 cells and MsNACs

## Footnotes

1 Schulze, E., 2021. "Premarket notice for integral tissue cultured poultry meat.". https://www.fda.gov/media/163262/download ; Good Meat Inc., 2022. “Dossier in support of the safety of GOOD meat cultured chicken as a human food ingredient.” Works Cited https://www.fda.gov/media/166346/download.

